# Evolved resistance against the type 6 secretion system is toxin specific

**DOI:** 10.64898/2026.03.17.712380

**Authors:** William Smith, Alejandro Tejada Arranz, Raveen Tank, Marek Basler, Michael Brockhurst

## Abstract

Across ecosystems, microbes face attacks by competitors armed with lethal weapons, including toxin-injecting Type 6 Secretion Systems (T6SSs). This in turn selects for T6SS resistance, reducing the future effectiveness of T6SS weaponry in competition. However, we have only a limited understanding of how resistance to T6SS attacks evolves *de novo*. A key challenge is that T6SS-armed bacteria are highly diverse in the number and type of toxins they inject, spanning multiple distinct modes of action. Do functionally distinct T6SS toxins select for different mechanisms of resistance? To address this, we combine genomics, resistance assays and fluorescence microscopy, to characterise evolved resistances against common T6SS toxin classes across spatial scales. We discovered that amidase and lipase toxins select for mutations in distinct sets of genes, resulting in toxin-specific resistance phenotypes and fitness costs. We also discovered resistance trade-offs: lipase-evolved *E. coli* became less vulnerable to lipase membrane damage, at the cost of increased susceptibility to lysis by amidase toxins. Finally, using single-gene knockout mutants from the Keio collection we confirmed that specific genes not previously linked to T6SS resistance, including inner membrane transporters, osmo-sensing systems and stress response pathways, conferred resistance to specific toxins while generating sensitivity to others. The specificity of resistance, and associated trade-offs observed, are likely to constrain *de novo* evolution of resistance against T6SS attackers armed with multiple functionally distinct toxins, helping to explain why T6SS systems are so widespread in nature.

## Introduction

Aggression is central to the ecology and evolution of microorganisms^1–3^. Killing other microbes can benefit bacteria both as means to feed on other cells^4^ (predation), and to prevent competitors from seizing valuable resources^2^ (interference competition). Bacteria in particular use a range of toxic weaponry to attack other microbes, enabling them to establish, expand and defend their ecological niches^5–7^. Significantly, such weaponry enables animal and plant pathogens to overcome colonisation resistance and invade established microbial communities on and within hosts^8,9^. A predictive understanding of microbial susceptibility to competitor attacks, therefore, would inform microbiome ecology and infectious disease^10–21^.

The Type 6 Secretion System (T6SS) is a ubiquitous weapon system found in 25% of gram-negative bacteria, including a variety of animal- and plant pathogens^22^. Among other roles, the T6SS mediates ecological competition within mammalian and insect microbiomes^8,23–25^ as well as in plant and soil ecosystems^26,27^. With a structure resembling the tail of a contractile bacteriophage, the T6SS functions by piercing neighbouring cells, translocating protein effector toxins^28^ that then damage rivals by targeting the cell wall, membranes, chromosome, and core metabolic functions^29–31^.This contractile mode of action gives the T6SS a short spatial range^32,33^, but also confers the unusual ability to deliver multiple toxic effectors in a single translocation event, confronting the target cell with a multitude of orthogonal stressors^15,34^.

The ubiquity of T6SS antagonism across ecosystems suggests that selective pressures to escape T6SS killing will be pervasive. Consistent with this, there is now extensive evidence that T6SS antagonism has driven the evolution of specific^35^ and generic^11,17,21^ defences against T6SS attack. Immunity proteins—the earliest known mechanism of T6SS protection— are central to how many bacteria escape T6SS antagonism^5,36,37^. Attackers express these proteins alongside toxins, enabling them to escape the effects of cognate toxins secreted by kin and competitors. Beyond classical immunity, genetic screen experiments have identified alternative mechanisms conferring specific or general resistance to T6SS attacks^11,35,38^. Known resistance mechanisms include individual-scale defences such as target modification^39^, removal of toxic effector products^35^, capsule production^10,11,20^ and specific and general stress responses^11,40^. There are also multiple mechanisms of collective resistance: populations of bacteria avoid attackers via biofilm^14,40^ or microcolony formation^19,41^, and via manipulation of colony spatial structure^42^. Intriguingly, these collective mechanisms appear to confer general (pan-) resistance against any combination of T6SS toxins, and these could emerge, in principle, via simple and accessible mutations (e.g. loss of repressor proteins^19^).

While multiple mechanisms of T6SS resistance are known, it remains ambiguous how defensive adaptations evolve during competition. This problem is made more complex by the multiplicity and functional diversity of toxins that T6SSs inject. When and how do defensive adaptations evolve *de novo* during competition via different toxin effectors? Do different effectors select for different kinds of defence, or do we just see pan-resistance evolve every time? To address these questions, we characterised resistance genotypes and phenotypes in *E. coli* strains experimentally evolved against different T6SS toxins^34^. Using whole genome sequencing, we show that toxins targeting the cell wall or membranes select for distinct sets of mutations, acting on a partially overlapping set of envelope biogenesis pathways (peptidoglycan editing, antigen, lipopolysaccharide and fatty acid biosynthesis). We develop colorimetric resistance assays to confirm that resistance phenotypes are primarily toxin specific at the population level, with occasional instances of cross resistance or collateral sensitivity. At the single-cell scale, we use fluorescence microscopy to show that lipase-evolved *E. coli* lineages become near-insensitive to lipase attacks, at the cost of increased vulnerability to amidase toxins – whereas amidase resistance is more subtle and potentially the product of a collective mechanism. Finally, using single-gene knockout strains from the Keio collection^43^, we confirm that loss-of-function mutations in *E. coli* can individually generate toxin-specific resistance, while also increasing susceptibility to other toxins. Our work suggests that the evolution of T6SS resistance is toxin specific and is constrained not just by resistance mutation costs, but by the narrow spectrum and underlying trade-offs between resistance mechanisms.

## Results

### Different T6SS effectors lead to non-overlapping sets of parallel mutations in *E. coli*

We began with a collection of 28 *E. coli* isolates^34^, drawn from populations experimentally evolved against *A. baylyi* T6SS attacker bacteria secreting cell-wall-targeting amidase effectors (Tae1), membrane-targeting lipase effectors (Tle1), both effectors together (Tae1+Tle1), or neither effector (T6SS− *Δhcp* control). We previously assayed these isolates for resistance to Tae1 and Tle1, revealing high levels of evolved resistance (up to 1,000x) in strains evolved against single-toxin effectors (Fig. 1A), but negligible resistance in control lines and lines evolved against the double-effector attacker^34^.

**Fig. 1:**
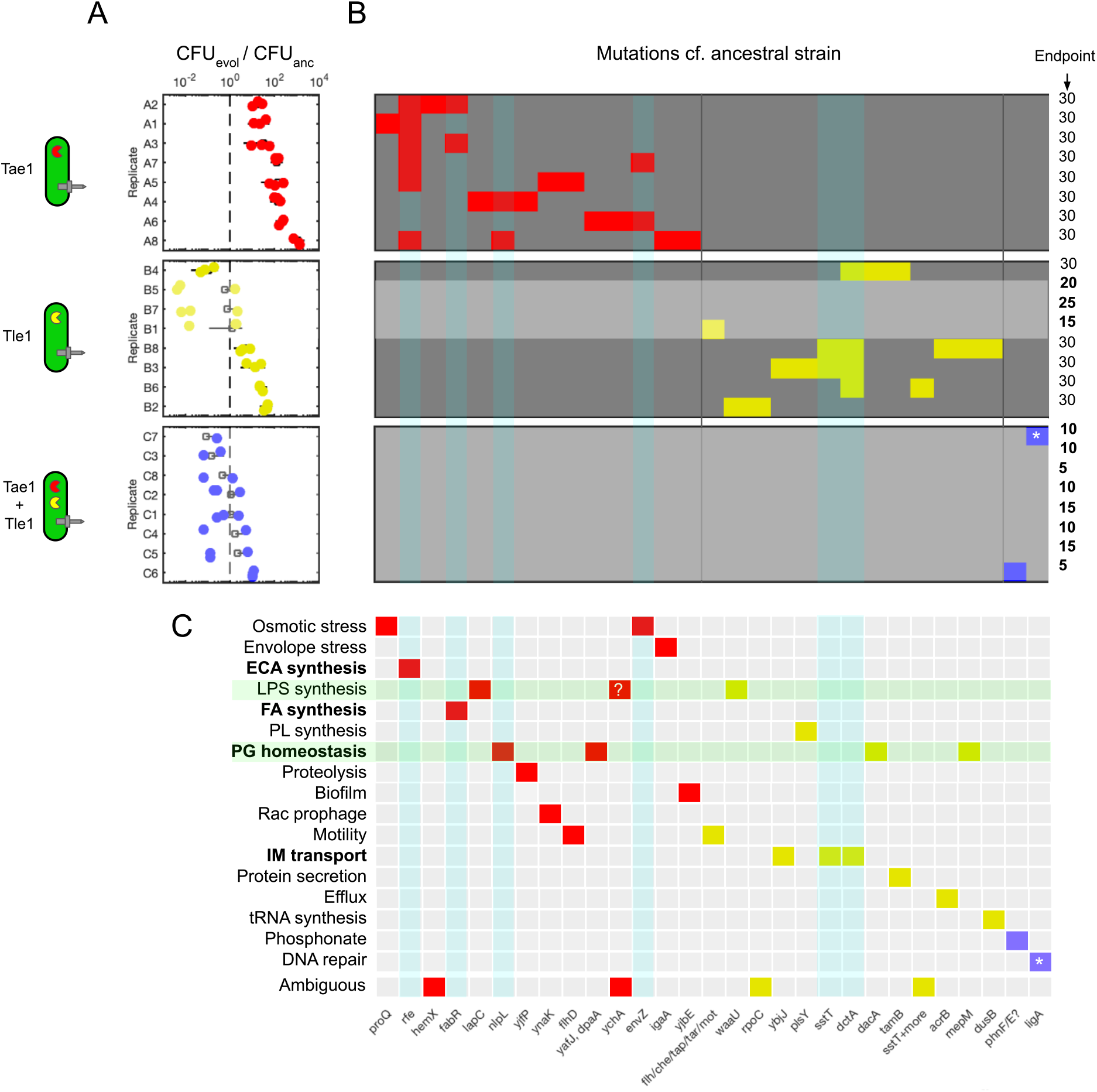
different T6SS effectors drive alternate resistance mutations. **(A)**: measurements of *E. coli* survival in competition with cognate attacker, relative to ancestral strain. N=3 pseudobiological replicates; dashed line represents ancestral survival against each competitor. Data replotted from previous publication^34^ **(B):** Presence/absence matrix indicating mutations in specific genes (horizontal axis) across evolved isolates (vertical axis); coloured by treatment type (red: amidase Tae1; yellow: lipase Tle1; blue: both effectors). Parallel mutations are highlighted with cyan overlay. End-point isolates drawn from populations that went extinct before the complete 30 transfers are a highlighted with a light grey overlay. **(C):** Presence/absence matrix associating mutations with gene ontology function terms collected via EcoCyc database, again coloured by treatment type. Functions altered in both amidase and lipase treatment groups are highlighted with green overlay; pathways for parallel mutations in (B) are highlighted in bold. *: synonymous (A514A) *ligA* mutation in strain C7. ?: IS1 mutation to uncharacterised gene *ychA* may affect downstream *kdsA* involved in lipopolysaccharide synthesis.

First, we wanted to know which genetic changes underpinned these resistance phenotypes, and so we carried out whole-genome sequencing (WGS) on all 28 evolved *E. coli* clones, identifying and annotating mutations using Breseq (See Methods). After removing preexisting mutations present in our ancestral *E. coli* strain, we identified mutations arising repeatedly across, and within, each of our T6SS treatment groups (Figs. 1B and S1). As is typical for the *E. coli* K-12 background, we saw a high frequency of mutations mediated by insertion sequences (ISs; 32 / 63 mutations), as well as substitutions and small indels. Across treatments, the most common mutation was a 1,199 bp deletion corresponding to loss of the IS5 Transposase *insH (*2-3 instances per treatment; 10 overall, Fig. S1A). Within the T6SS− control group, we saw repeated loss of flagellar motility and chemotaxis genes, previously associated with *E. coli* adaption to liquid media^44,45^ (Fig. S1E). Because they occur irrespective of or absent T6SS killing, both mutation types likely represent adaptation to lab culture conditions rather than to T6SS selection, and so we excluded these mutations from further analyses.

After filtering mutations, we saw that amidase and lipase selection led to distinct, non-overlapping sets of non-synonymous mutations (Fig. 1B), suggesting strong, toxin-specific selective pressures. Parallel evolving loci unique to amidase selection included frequent disruption of the *rfe* (aka *wecA*) gene or its promoter region (6/8 lines), substitutions in *fabR* (2/8), substitutions or nonsense mutations in *nlpI* (2/8) and substitutions in *envZ*. Strikingly, these mutations affect not only the cell wall (the direct target of the amidase) but various cell envelope components (*nlpI*: peptidoglycan biosynthesis; *rfe*: biosynthesis of enterobacterial common antigen, ECA; *fabR*: fatty acid biosynthesis regulator; *envZ*, osmosensing regulation). Meanwhile, parallel mutations unique to lipase selection included mutations to *sstT* (3/8) and to *dctA* (4/8), both of which encode inner membrane transporters. Additionally, we saw a clear relationship between number of mutations and extinction: *E. coli* lines that went extinct before the end of the 30 day passaging experiment (Tle1: B1,5 and 7; C1-8; light grey rows in Fig. 1B) showed fewer or no mutations, consistent with a failure to adapt to T6SS attack resulting in extinction (Fig. 1B).

Unexpectedly, we saw few instances of mutations previously associated with resistance to other T6SS toxins (exceptions: *lapC* involved in LPS synthesis^17^ in A4; *igaA* repressor of the Rcs envelope stress response system^11^ in A8). The majority of our observed mutations are, to our knowledge, members of pathways not previously associated with T6SS resistance. We were particularly surprised to see peptidoglycan maintenance gene mutations (e.g. *mepM, dacA*) appear in multiple lipase-treated lines as well as in multiple amidase-treated lines (*nlpI, dpaA*), albeit targeting distinct genes per toxin. Previous work has shown that modifications to membrane lipopolysaccharide biosynthesis can confer resistance to amidase toxins^16^; our observations suggest that, similarly, alterations to cell wall synthesis could protect against membrane-targeting lipase effectors.

To further explore the types of biological processes affected by mutations, we examined the gene ontology (GO) terms associated with each mutated gene using the databases EcoCyc^46^ and UniProt^47^, using these to classify each gene’s broad function (Fig. 1C). Amidase-associated mutations tended to affect genes associated with cell envelope stress or maintenance, whereas lipase-associated mutations mostly commonly involved inner membrane transport. Unlike at the gene level, we saw partial overlap in the cellular processes affected by mutations, with both treatments selecting for modifications to PG homeostasis, and LPS and FA synthesis pathways. This limited functional overlap suggests that, while the two toxins select for mutations in distinct sets of genetic loci, these may converge on similar cellular processes and pathways.

In summary, our WGS data show that different T6SS toxin types select for parallel and treatment-specific sets of mutations, many of which have not previously been associated with T6SS resistance, and that these mutations facilitated evolutionary rescue of surviving cell lines. These findings suggest that, resistance evolved in our experiment appears to be toxin-specific rather than generic, contrasting recent studies identifying pan-resistance mechanisms in gene deletion screens^10,14,19^.

### Population- and cell-scale assays reveal toxin-specific resistance phenotypes and trade-offs

Protection against T6SS attacks can arise via both individual- and collective resistance mechanisms^14,15,19^. Given the effector-specific mutations identified above, we wanted to compare the resistance phenotypes arising in amidase- and lipase-evolved *E. coli*, both at the level of individual cells and as cell collectives. We therefore developed assays to phenotype amidase- and lipase resistance at both spatial scales (Fig. 2, Methods). Inspired by previous work^48^, our colorimetric population-scale assay (Figs. 2A-C, S2 and S3) uses the chromogenic marker 5-bromo-4-chloro-3-indolyl-D-galactopyranoside (“XGal”): live *E. coli* bacteria convert XGal into a blue-green pigment, providing spatial maps of survival (pigment intensity) across ∼5 mm colonies, and semi-quantitative indicator of population-scale resistance (Fig. S2A, B). Meanwhile, our single-cell assay uses confocal microscopy and a cell permeabilization reporter dye (SYTOX blue) to intensively track rates of *E. coli* killing per contact with *A. baylyi* attackers^49^ (Figs. 2D-F and S4). Both assays were performed using high density *A. baylyi / E. coli* mixtures (1:1 mix at OD_600_ = 10), providing a robust test of resistance under high T6SS attack pressure.

**Fig. 2:**
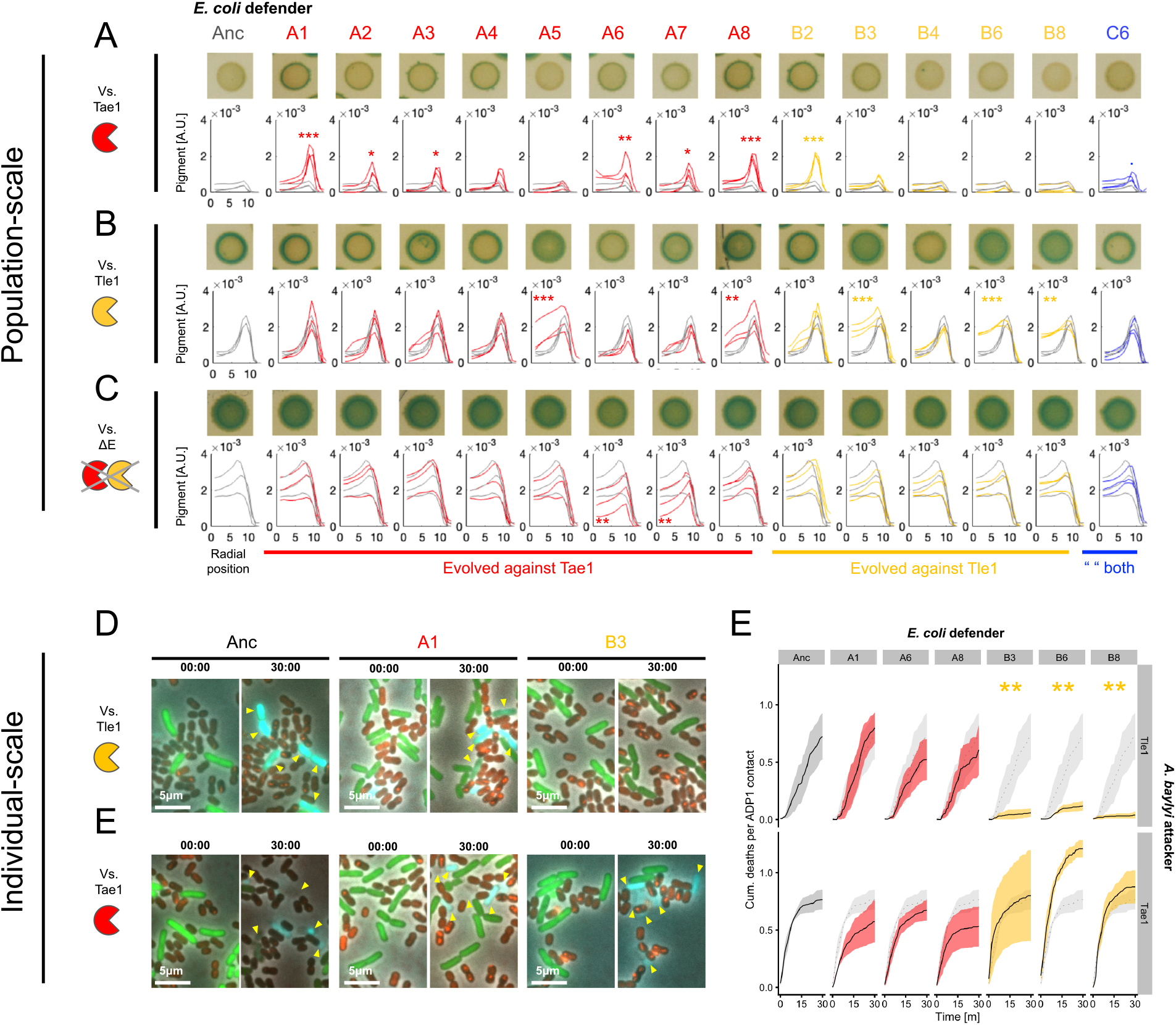
Population- and cell-scale killing assays implicate toxin-specific resistance mechanisms operating at alternate spatial scales. **(A)** *E. coli* defenders (columns) in 16h 1:1 co-culture with Tae1 amidase-armed *A. baylyi* T6SS attackers, plated on chromogenic XGal agar. Green-blue pigmentation highlights colony locations where *E. coli* survive. Representative colonies (biological replicate 1) are shown above plots of radial pigment intensity traces (red: Tae1-evolved; yellow: Tle1-evolved; blue: Tae1+Tle1-evolved), with XGal traces from the ancestral strain plotted (grey lines) for reference. Asterisks indicate traces with XGal significantly greater than that for the ancestral strain at the colony edge (***: p<0.001; **: p < 0.01; *: p<0.05); statistical tests using linear regression (R: “lm” method using Dunnett’s contrast for multiple comparisons). **(B)** as (A) but with Tle1 lipase-armed *A. baylyi* T6SS attackers; asterisks mark isolates with increased survival in the colony centre. **(C)** as (B) but with effectorless (ΔE) *A. baylyi* control strain. **(D)** Representative snapshots from 30m timelapse microscopy, comparing responses of Anc(estral), A1 (Tae-evolved) and B3 (Tle1-evolved) *E. coli* isolates to lipase Tle1. Image timestamps and scale bars as shown; eGFP+ *E. coli* cells, T6SS foci (*vipA::mCherry* fusion) and SYTOX permeability stain are respectively coloured green, red and cyan. Yellow arrows highlight individual cell death death events. **(E)** as (D), but against the amidase Tae1. **(F)** Quantification of single-cell killing dynamics using *E. coli* deaths (SYTOX+ events), normalised by number of *E. coli* - *A. baylyi* contacts, for select *E. coli* isolates (columns) vs. single-effector attackers (rows). Lines and shaded areas (coloured according to evolutionary treatment as in A) correspond to mean +/-standard error; the ancestral traces are included in each panel for comparison. Asterisks indicate killing traces with AUC significantly different from the ancestral strain (**: p<0.01); statistical tests using linear regression (R: “lm” method using Dunnett’s contrast for multiple comparisons). N=3 biological replicates used for each strain pairing.

We conducted colorimetric assays on all *E. coli* isolates that survived the full 30-transfer time evolution experiment, as these initially showed the strongest resistance phenotypes. We also included C6 as this was the only isolate to show apparent resistance to the double-effector attacker. By measuring pigment intensity as a function of colony radial position (representative colonies and traces shown Fig. 2A-C), we confirmed that evolved *E. coli* have high levels of resistance to focal effectors: excepting A5 (which, curiously, displayed Tle1 resistance), all Tae1-evolved isolates survived Tae1 treatment better than the ancestor (Fig. 2A), with survival improving specifically at the colony edge (Fig. S2C). Conversely, all Tle1-evolved isolates (except B4, which was also non-resistant in our original CFU assay) survived Tle1 treatment better than the ancestor, with survival improved in the colony centre (Figs. 2B, S2C).

While strains showed high levels of resistance against the toxins they had previously evolved against, only isolate A8 showed resistance to both toxins (Fig. 2A,B and S2C). Additional colorimetric assays showed that A8’s cross-resistance phenotype is also limited to specific effectors (Fig. S3): A8 behaved identically to the Ancestor against WT *A. baylyi* ADP1 (secreting effectors Tae1, Tle1 and Tse2, Fig. S3A,B) and WT *Vibrio cholerae* 2740-80 (secreting VgrG3+TseH+TseL+VasX, Fig. S3C,D), with some weak resistance against *V. cholerae* lacking active pore-forming toxin VasX. Overall, these results confirm that the results of the CFU assay (Fig. 1A) are robust with respect to very high cell densities, but that – consistent with the toxin-specific mutation sets observed in Fig. 1 – evolved resistances are primarily toxin-specific, granting protection against the focal toxin but not to other toxins in general.

Given that the majority of our evolved *E. coli* isolates have mutations potentially affecting the cell surface (e.g. *wecA* nonsense mutations leading to antigen loss^50^) we considered that resistance might involve mechanisms of cell aggregation (e.g. formation of protective microcolonies via increased cell adhesion^19,41^) or altered deposition (e.g. via the “coffee ring” effect^51^, leading to reduced contact with attackers^42^), potentially explaining why different isolates showed survival in different colony locations. Across treatments however, we saw no evidence of cell aggregation or microcolony formation, which would be expected to produce granularity in XGal pigmentation. Similarly, assays performed with effectorless *A. baylyi* attackers (Fig. 2C), showed that, absent T6SS intoxication, strains have comparable edge and centre pigmentation to the ancestor, with weaker growth seen in isolates A6 and A7 (Fig. S2C). We therefore rejected the hypothesis that evolved resistance emerged via microcolony formation, or innate biophysical segregation of strains, as reported previously.

Using fluorescence microscopy, we then examined whether T6SS resistance or cross-resistance would manifest in individual *E. coli* cells. We selected a subset of three *E. coli* isolates per evolutionary treatment (those with the strongest population-scale resistance phenotypes vs. their cognate attacker: A1, A6, A8 and B3, B6, B8). As previously^49^, interaction with amidase-secreting ADP1 led to explosive cell lysis and blebbing of the ancestral *E. coli* strain, whereas lipase Tle1 led to slow cell permeabilization (Figs. 2D,E and S4A).

In contrast to the ancestor, lipase-evolved *E. coli* showed significantly reduced permeabilization against lipase-armed attackers (example: Figs. 2D and S4C), indicating that individual cells are strongly resistant to this toxin (Figs. 2F and S4C), consistent with their strong population-scale resistances. Strikingly however, this resistance came at the cost of faster lysis against Tae1-armed attackers (Figs. 2D,F and S4C), indicating a resistance trade-off arising from the *dctA* and/or *sstT* mutations these isolates share. Our population-scale assay did not identify this trade-off, potentially because the Tae1 treatment (Fig. 2A) leads to so little *E. coli* survival after 16h that increased Tae1 sensitivity could not be detected. Meanwhile, compared with the ancestral strain, isolates A1, A6 and A8 showed only a marginal reduction in killing by Tae1 (Fig. 2F). Given the strong population-scale resistances observed here, this result was unexpected and may implicate a collective resistance mechanism (i.e. where bacteria are resistant as a population but not as individuals)^14,15^. These differences between treatment groups are consistent both with the toxin-specific genetic modifications we see in these isolates (Fig. 1), and the toxin-specific resistance phenotypes exhibited in the population-scale assay (Fig. 2)

In summary, mutations found in lipase-evolved isolates result in strong, individual-level resistance at the cost of amidase sensitivity, whereas mutations found in amidase-evolved isolates—while producing strong resistance at the population level—generate more subtle differences in killing at the individual scale, implicating collective resistance mechanisms to reconcile these findings.

### Amidase resistance confers greater fitness costs than lipase resistance

T6SS resistance has been associated with reproductive fitness costs that increase with the number of mutations present^17^. Paralleling resistance costs to other antimicrobials^15^, these are expected to constrain the evolution of resistance by slowing selective sweeps for resistance, and selecting for resistance loss in the absence of T6SS aggression. Given that they impact multiple highly conserved cellular processes (particularly lipopolysaccharide and peptidoglycan synthesis, Fig. 1), we hypothesised that our amidase- and lipase-resistant mutants might suffer similar fitness costs.

To test this, we performed two assays to estimate fitness costs in evolved mutants, compared with the ancestral *E. coli* strain (Fig. 3). We measured mutant growth in broth monocultures over 24h (Fig. 3A). In parallel, we performed pairwise competition between evolved mutants (eGFP-labelled) and the ancestral *E. coli* strain (mCherry2-labelled), using flow cytometry to measure changes in the mutant : ancestor ratio (Figs. 3B and S5) during co-culture (see Methods). We included the latter assay because i) resistant strains with altered cell envelopes may have altered light scattering properties, such that optical density becomes a misleading proxy for cell density, and ii) monoculture growth assays do not capture interactions between the mutant and its ancestor: e.g. impaired nutrient update that is only apparent when a more efficient consumer is present. Both assays showed that evolved *E. coli* strains have a range of fitness impairments, whose magnitude depends on the toxin treatment used during experimental evolution. Control lines (evolved without T6SS antagonism) showed slower growth compared with the ancestor in monoculture (Fig. 3A), but with the exception of isolate D4, this did not result in lower competitiveness in coculture (Fig. 3B).

**Fig. 3:**
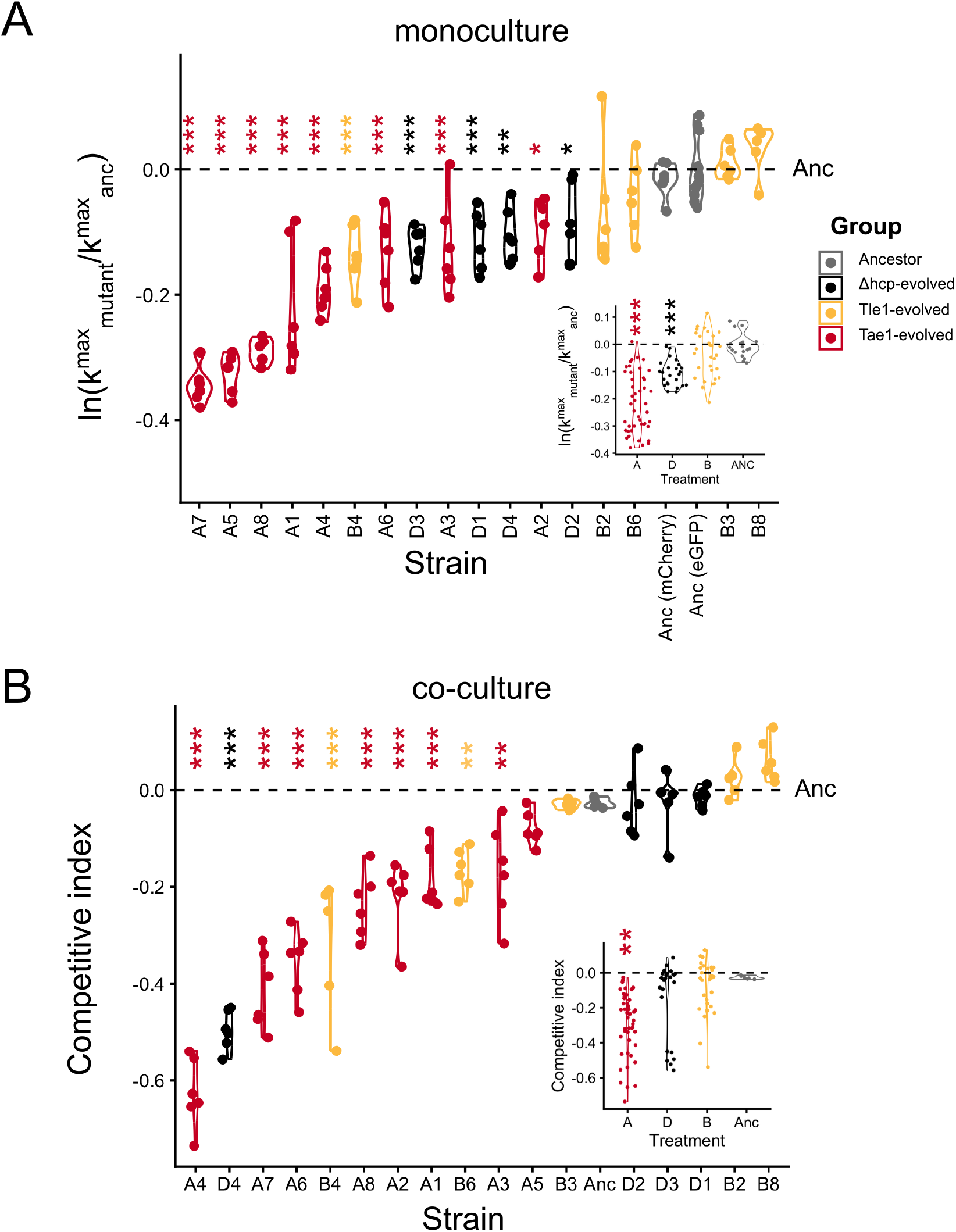
Mono- and co-culture fitness assays show amidase resistance is costly while phospholipase resistance is cheap. **(A)** Violin plot showing stains’ maximum growth rates (k_max_) in liquid monocultures, plotted relative to the eGFP-labelled ancestral strain. Inset: variation in scaled maximum growth rates between evolutionary treatments. Dashed lines indicates the mean max growth rate of the ancestral strain, N=6 pseudobiological replicates for each strain monoculture. **(B)** Mutant competitive index showing eGFP-labeled mutant fitness relative to mCherry-labelled ancestral strain in 1:1 liquid co-cultures, measured using flow cytometry. Dashed line indicates the competitive index of the ancestral strain. Inset: variation in competitive index between evolutionary treatments. Data points and violin curves are coloured according to evolutionary treatment group (see legend). Asterisks denote significant deviations from ancestor: (***: p<0.001; **: p < 0.01; *: p<0.05); statistical tests use linear regression (R: “lm” method using Dunnett’s contrast for multiple comparisons); N=6 pseudobiological replicates used for each co-culture.

Significantly, in both mono- and co-culture fitness assays and Fig. 3C, respectively), amidase-evolved isolates showed substantially reduced max growth rate (Fig. 3A) and reduced growth (Fig. 3B) relative to the ancestral strain, despite them having fewer mutations on average than phospholipase-evolved isolates (3.125 for Tae1 *cf*. 3.6 per strain for Tle1). In contrast, Tle1-evolved isolates showed negligible changes in mean fitness in both mono- and co-culture assays. A notable exception was strain B4, which shows both minimal resistance (Figs. 1A and 2A, B) and high fitness costs (Fig. 3), leading to the question of how it evolved in our serial transfer experiment. We hypothesise that the narrow bottleneck of the (more lethal) Tle1 treatment resulted in chance fixation of a deleterious mutation during passaging, which we posit to be *dacA* and/or *tamB* mutations unique to this line. *dacA* mutations generate severe morphological defects in cells^52^ (confirmed using microscopy, Fig. S4E) and this could negatively impact both fitness and T6SS sensitivity. Overall, these results show that fitness costs associated with T6SS resistance are i) variable (and in some cases essentially undetectable in liquid culture) and ii) toxin-specific, with amidase resistance generally being costlier.

### Competition assays with Keio isolates identify specific loss-of-function mutations leading to toxin resistance and collateral sensitivity

Having explored the specificity of resistance genotypes and phenotypes, we sought finally to associate toxin-specific resistances with individual mutations in *E. coli*. To do this, we conducted additional resistance tests using isolates from the Keio collection^43^, a set of 3,985 *E. coli* isolates with individual non-essential gene deletions. Keio isolates have previously been used both to screen for genes involved in T6SS susceptibility^38^ and to estimate the effects of mutations occurring in the K-12 MG1655 (ancestral) background^44,53^, this being similar to the Keio parent strain K-12 BW25113. While only a subset of the mutations described above are likely to result in complete loss of gene function (e.g. via deletion, frameshift or nonsense mutations), we reasoned that deletion of the corresponding gene might still alter T6SS resistance via perturbation of the underlying pathway. Because Keio isolates intentionally lack a functional *lacZ* gene, we could not use our colorimetric XGal assay here, and so we used a standard (CFU-based) competition assay^14,32,34,49^.

We performed two competition assays (Methods) using Keio isolates (Fig. 4), initially prioritising the most parallel mutations seen in each single-effector treatment group (*rfe*/*wecA* for Tae1; *dctA* for Tle1; Fig. 4A). We used the Keio *ΔlacY* strain as a control, reasoning that the *lac* operon is already disrupted in this background and so loss of *lacY* (encoding the lactose symporter) would have no effect on T6SS susceptibility. We also included the original MG1655 ancestral strain to compare susceptibility differences between these two backgrounds. We were surprised to find that loss of the *rfe* gene had no effect on *E. coli* survival in competition with the amidase-armed attacker (Fig. 4A), despite this being the most common mutation seen in amidase-evolved *E. coli* (6/8 lines). Loss of *dctA* also had no effect against amidase attacks, but did grant significant resistance to attackers armed with the lipase. We noted that, against both attacker strains, the ancestral MG1655 strain had lower survival than the Keio *ΔlacY* strain, an effect which might arise from genetic differences between the two K-12 backgrounds. For instance, the Keio parent has a predicted loss-of-function mutation in its *fabR* gene^54^, which we also saw mutated repeatedly in Tae1-treated cell lines.

**Fig. 4.**
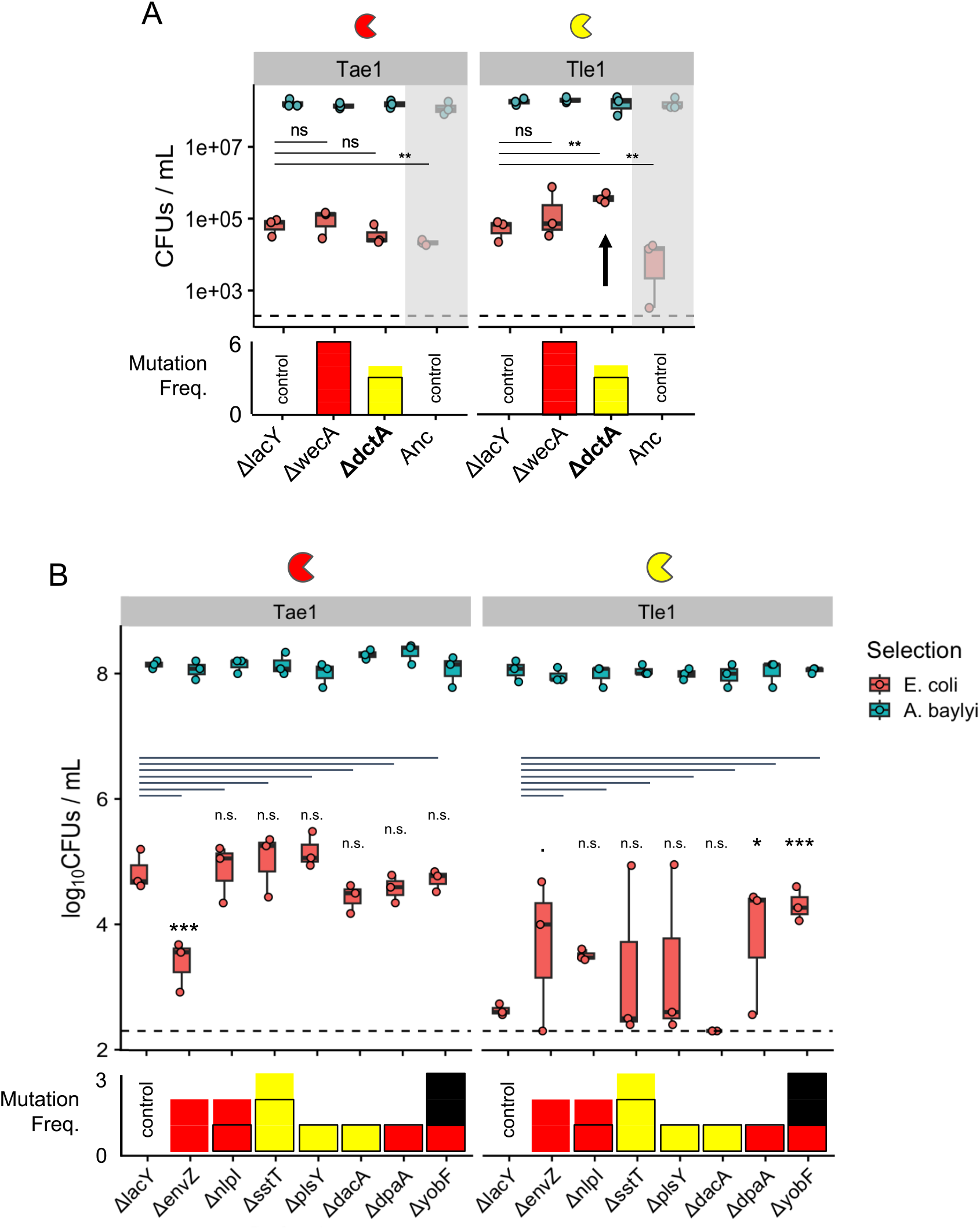
T6SS resistance assays using Keio single-gene knockout strains identify new resistance determinants. **(A):** initial competition assays quantifying changes in *E. coli* sensitivity to Tae1 (left) or Tle1 (right) arising from single gene deletions, targeting the most common parallel mutations identified in each single-effector treatment: *ΔwecA* (*Δrfe*) and *ΔdctA* for Tae1, Tle1 respectively. In the upper boxplot, bold lines, boxes and whiskers correspond respectively to data means, interquartile ranges and ranges. The neutral *ΔlacY* knockout is the control for the Keio strain background; we also included the Anc control for comparison. **(B)** as (A) but showing subsequent tests examining remaining parallel and selected singleton mutations. In A and B, the lower bar graph plots the number of parallel mutations to each gene (bar colours indicate the treatment group(s) in which these were identified: red=Tae1, yellow=Tle1; black=T6SS-control); the black outlines indicate the proportion of mutations likely to lead to loss of gene function). Statistical tests use linear regression with replicate as fixed factor (R: “lm” method using Dunnett’s contrast for multiple comparisons); significance codes ***: p<0.001, **: p<0.01, *: p<0.05, :? p<0.1, n.s.: p≥0.1. N=3 full biological replicates per strain combination, each the average of 3 technical replicate platings of the recovered *E. coli / A. baylyi* cell mixtures post competition.

We then repeated our assay to examine amidase- or lipase sensitivity changes arising from other gene loss events (Fig. 4B). We again prioritised testing of knockouts where the deleted gene saw parallel mutations during experimental evolution (*ΔenvZ, ΔnlpI* against amidase attacks, *Δsst* against lipase attacks, see Fig. 1B). We also included four singleton mutations with predicted functions of interest (*ΔplsY, ΔdacA, ΔdpaA* and *ΔyobF*). Some knockouts could not be included in this assay, either because of preexisting loss-of-function mutations in the Keio parent (*ΔfabR*, see above), or because mutations occurred in an essential gene (*ΔigaA, ΔwaaU*). This assay showed that, intriguingly, *envZ* gene loss results in ∼100x increase in sensitivity to the amidase, but variable resistance to the lipase, demonstrating that collateral effector sensitivity can emerge via a single loss-of-function mutation. *ΔnlpI, ΔdpaA* and *ΔyobF* mutations had no effect against Tae1 but appeared to increase resistance against Tle1 (very significantly in the case of *ΔyobF*, which is consistent with the unexpected lipase resistance in the *yobF* mutant A5). By contrast, *Δsst, ΔplsY* and *ΔdacA* knockouts had no significant effect on T6SS resistance to either effector (however, *ΔlacY* survival vs. Tle1 was close to the limit of detection, meaning that potentially sensitising effects of *dacA* loss could be masked). This could suggest that these mutations only affect phenotype in combination with other mutations (i.e. positive epistasis).

Overall, our knockout strain tests show we can recreate some, though not all, resistance phenotypes arising from specific mutations in evolved strains. Mirroring the results of our colorimetric and microscopy assays, we saw toxin-specific resistance in single-gene knockouts (IM transporter *ΔdctA*, stress response protein *ΔyobF*, peptidoglycan amidase *ΔdpaA*); in the case of *ΔenvZ*, lipase resistance also came at the cost of significant amidase resistance.

## Discussion

T6SS antagonism is an important and pervasive component of many microbial communities and ecosystems, suggesting widespread selective pressures for T6SS-defensive adaptation in microbes. Previous work has used genetic screens to identify bacterial genes involved in generic and specific resistance to T6SS attack^11,16,19,21,35,38^. Here, we show that *E. coli* bacteria spontaneously evolve novel and toxin-specific resistance mechanisms against two common T6SS effector classes, amidases and lipases. We analysed the genotypes and phenotypes of *E. coli* that had been experimentally evolved to resist two T6SS effectors, amidase Tae1 and lipase Tle1, both secreted by *A. baylyi* ADP1^49^. This revealed that alternate sets of resistance mutations occur repeatedly in these two treatment groups, and that amidase resistance likely involves alterations to envelope biogenesis pathways and osmosensing systems. Meanwhile, lipase resistance was associated with modification or loss of inner membrane transporters *sstT* and *dctA*.

Using chromogenic XGal agar to map *E. coli* survival at a population scale, we confirmed strong toxin-specific resistance (and occasional cross-resistance) phenotypes, but ruled out pan-resistance mechanisms via cell aggregation or escape to the colony edge. We also examined resistance phenotypes at the individual cell level using fluorescence microscopy, demonstrating a resistance trade-off: strong individual-scale lipase insensitivity came at the cost of increased vulnerability to the amidase toxin Tae1. In contrast, amidase-evolved isolates showed only marginal resistance to the amidase at the single-cell level; the weakness of this effect compared with the strong resistance apparent in population-scale assays (Figs. 1A and 2A) suggest that a significant component of resistant arises via an unknown collective mechanism, other than aggregation or escape to the colony edge.

Using monoculture and co-culture growth assays, we also identified asymmetries in resistance fitness costs: while generally stronger in effect, amidase resistance was more costly compared with lipase resistance. These single-effector resistance costs are likely to constrain the evolution of multi-toxin resistance: if, as in our experiments, pan-resistance mutations are rare or inaccessible, multi-resistance would require the accumulation of costly mutations against each effector. Assuming additive epistasis, resistance to a large set of effectors would then be at least as costly as the costliest effector, even if resistance to some effectors is low-cost. We also note that our (single-effector) resistance costs are substantially lower than those reported by MacGillivray and colleagues (multi-effector selection)^17^, pointing to single-effector resistance as being generally cheaper than multi-effector resistance. High cumulative costs could also explain why, although clearly feasible, multi-resistance was much rarer evolutionary outcome than single-toxin resistance.

Finally, we used *E. coli* gene deletion mutants to show that loss of *dctA, envZ, dacA* and *yobF* genes all confer significant but toxin-specific resistance phenotypes, offering protection against one toxin, but no protection (or enhanced susceptibility in the case of *envZ*) against others. Mirroring collateral antibiotic sensitivities seen by others^17,21^, we hypothesise that these effector sensitivity trade-offs produce additional constraints that further limit T6SS resistance evolution, e.g. by requiring bacteria to accumulate multiple actively-conflicting mutations to achieve resistance to multi-toxin T6SS attacks (i.e. antagonistic pleiotropy).

To our knowledge, ours is the first study of experimental resistance evolution vs. specific T6SS effectors. This approach complements previous work based on knockout screening using TnSeq^35,55^, CRISPRi^16,21^ or gene knockout libraries^19,38^. Screening approaches are valuable because they test the effects of the broadest possible range of mutations, examining the effects of loss or reduction of any target gene. Meanwhile, experimental evolution tests what evolves in practice during repeated competition, incorporating the complexities of fluctuating selection and population demographics, mutational supply and biases, genetic drift, interactions between mutants (clonal interference, fitness costs) and mutations (epistasis, pleiotropy). Reassuringly, the two approaches converge on the same answers in some cases: we saw multiple mutations affecting pathways (e.g. *igaA* affecting the Rcs stress response; *LapC, WaaU* and potentially *kdsA* affecting LPS synthesis) identified as mediating resistance to single or multiple toxins by genetic screens. However, the limited overlap between our resistance mutations and those identified previously against other T6SS effector sets further underscores the specificity of T6SS resistance.

Our study also identifies T6SS resistance routes not previously identified, including mutations to peptidoglycan maintenance pathways associated with amidase (via *nlpI, yafK*) or lipase selection (*dacA, mepM*). We note that several of the resistance genes identified are regulated by the cAMP receptor protein (CRP), whose inactivation by glucose was previously observed to confer resistance to *V. cholerae* T6SS attacks in *E. coli* via an unknown mechanism^56^. Suppression of *dctA* (outcome of *dctA* loss-of-function mutations) and *ompF* (shown previously to be the outcome of our envZ^P41L^ mutation^57^) are both downstream effects of CRP inactivation, suggesting that removal of these susceptibility factors may contribute to the resistance phenotypes observed by Crisan *et al*.

Pan-resistance mechanisms, such as biofilm or capsule formation, are clearly feasible^10,14,19^ yet did not evolve in our study. A possible exception is isolate A8, which showed an IS1 mutation in the promoter region of gene *yjbE*, which is potentially involved production of an extracellular polysaccharide^58^. A8 also exhibited limited cross-resistance to other amidase and lipase toxins secreted by the *V. cholerae μvasX* strain, and while it showed no visible EPS production in short timescale single-cell analyses, it could accumulate an EPS / biofilm barrier over longer timescales. Overall, the circumstances which select optimally for collective resistance mechanisms, like EPS production, remain poorly understood and it may be that our experimental evolution protocol excluded collective resistance phenotypes, by running selection stages at high cell density and attacker frequency. Exploring whether there are specific conditions favouring collective or pan-resistance would be an interesting avenue for future research. Going forward, it will also be important to check whether evolved resistances vs. other T6SS effector classes also have similarly high specificities to those we report here.

Overall, the mutational constraints we identified help to explain why multi-toxin resistance is comparatively difficult to evolve compared with single-toxin resistance^34^. Multi-toxin attacks are not only more lethal^30,34^, but also require the accumulation of multiple, potentially conflicting mutations before defenders gain multi-toxin resistance, making evolutionary rescue less likely than for single-toxin stressors. Natural selection would then be predicted to favour attackers whose weaponry remains robust to competitor resistance evolution (“biological robustness” with respect to competition^59,60^), and this in turn could explain why many T6SS-armed bacteria secrete multiple effectors with distinct mechanisms of action.

## Conclusion

Understanding microbial susceptibility to aggression is important both technologically (e.g. for designing microbial consortia with stable interaction networks), and ecologically (e.g. for predicting the susceptibility of communities to attack and invasion by pathogens). Our work shows that resistance to different toxin types requires alternate sets of mutations, and in some cases these occur through opposing alterations to a common set of pathways and homeostatic systems. This suggests that the evolution of T6SS resistance is constrained not only by resistance mutation costs, but by underlying molecular trade-offs in resistance mechanisms, helping us to understand and predict the outcomes of competition and evolutionary arms races.

## Methods

### Bacterial strains and cultures

Table S1 shows the bacterial strains used in this study. *E. coli* and *V. cholerae* strains were grown aerobically in 5 or 3mL of lysogeny broth (LB) at 37°c and 180 or 200 rpm. *A. baylyi* strains were grown similarly but at 30°c.

### Genome sequencing and genomic analysis

#### Whole-genome sequencing

To sequence the genomes of ancestral and evolved *E. coli* bacteria, we used the MicrobesNG standard sequencing service (MicrobesNG, Birmingham UK, https://microbesng.com). Exponential phase cultures of ancestral and evolved *E. coli* clones were prepared (1:50 dilution of overnight culture into 10mL fresh LB, incubation at 37°c, 180 rpm shaking for ∼3h). After reaching OD_600_∼1, we used centrifugation (20,000 g, 4 mins) to concentrate 1mL samples to OD_600_ ∼10.0, giving a total of ∼2.5×10^9^ bacterial cells. Bacteria were pelleted and resuspended in 500 uL DNA/RNA Shield Inactivation Buffer (Zymo research). Following shipping, DNA was extracted from cell lysates and purified using a NexteraXT library preparation kit, before being subjected to Illumina DNA sequencing (2×250 bp paired-end reads with minimum 30x coverage). Reads were trimmed (Trimmomatic v0.3, Q15 cutoff) and aligned to the reference using the Burrow-Wheeler Aligner (Maximal Exact Match algorithm).

#### Variant calling

To identify mutations arising in evolved *E. coli* isolates, we used breseq^61^ (v0.37.1, with bowtie v2.4.5) to align reads with an annotated *E. coli* MG1655 reference genome (genBank: U00096.3), using consensus mode and a coverage limit of 100. Our ancestral strain differed from the reference at 9 mutations (of which 5 were intergenic and 1 synonymous); we excluded preexisting mutations from analysis of evolved clone.

### Population-scale resistance assays

#### Colorimetric assay of population-scale resistance

We prepared exponential-phase cultures of *E. coli* and *A. baylyi* by diluting cultures grown overnight (∼18h) into 3 mL fresh LB (1:300 dilution for E. coli; 1:100 dilution for *A. baylyi*), before incubating for 3 h at 37°c and 200 rpm. Once exponential cultures reached an OD_600_ of >1 (*A. baylyi*) or >1.5 (*E. coli*) we harvested 1mL of each culture and concentrated it to OD_600_ = 10.0 using centrifugation and resuspension in fresh LB. All combinations of *A. baylyi* and *E. coli* cultures were mixed in a 1:1 ratio (5 μL of each suspension), and 5 μL of the resulting mix was spotted on 1.5% w/v LB agar plates supplemented with 40 μg/mL 5-bromo-4-chloro-3-indolyl-β-D-galactopyranoside (“XGal”) chromogenic substrate and pre-dried for 1h next to the Bunsen burner. Once the drops were dry (∼5 min), the plates were lidded, inverted and incubated at 30°c for 64h. After 16 and 64h, images were taken of each colony on a transilluminator using a digital camera. We used 3 biological replicates per strain combination.

#### Image processing to measure XGal pigmentation

To quantify *E. coli* survival at different locations within colonies, colony images (as individual .tif files, 300 x 300 px) were analysed using custom Matlab code. To quantify XGal pigmentation, we used a colormap whose extrema are 1) the edge of ancestral *E. coli* + ADP1 ΔE colonies (corresponding to max *E. coli* survival, max. XGal signal) and 2) the centre of ancestral *E. coli* + ADP1 Tae1 colonies (corresponding to min *E. coli* survival, min. XGal signal). We used Matlab’s 1-d interpolation function “interp1” to generate a 100-color linear colormap between these extrema. To account for lighting differences between plates, we generated an independent colormap for each biological replicate. We found that colony widths and centroids varied naturally within each image, requiring that images be aligned before comparing pigment signal intensity. To do this, we used a circular Hough transform (Matlab “imfindcircles” with radius range 80, 200 px, sensitivity=0.97) to automatically compute the centroid (x,y) and radius (r) of each colony image (as an inverted greyscale image). Fit quality was verified by eye for each colony and succeeded for 98.5% (n=133 / 135) of spots; for the remaining 2 spots we specified the colony centre and radius manually. Following colony segmentation, we used our colormaps to convert each RGB image to an array of colour indices (Matlab “rgb2ind” using the above pigment colormaps), ranging from 0 (minimum pigment) to 100 (maximum pigment). We divided each spot image into a set of concentric annuluses (as binary masks), each with a thickness of 0.1 r and centroid (x,y). To ensure coverage of the entire spot, we used a total of 13 annuluses (i.e. 0-130% of r). We then computed the per-pixel pigment intensity within each annulus as the sum of colour indices / 100. This procedure allows us to map XGal pigmentation, and by proxy *E. coli* survival, as a function of radial coordinate within each spotted mixture. To compare survival at different colony positions, we measured the average per-pixel pigment intensity within annuluses 30-40% of r (colony centre) and 90-100% of r (colony edge; see Fig. S2B, C).

### Fluorescence microscopy and image analysis

#### Time lapse microscopy

We grew overnight cultures of fluorescently labelled ADP1 attacker (Tae1- and Tle1-secreting VipA-mCherry fusions) and *E. coli* defender (constitutive eGFP tag, A1, A6, A8; B3, B6, B8) strains by inoculating 3 mL LB broth cultures with single colonies, and incubating overnight (37°c, 200RPM shaking). On the day of the experiment, we diluted overnight cultures 1:100 into 3 mL fresh media, growing for 3 h until cultures reached exponential phase (OD_600_ ∼0.8-1.2). Cultures were then concentrated to OD_600_=20, and mixed 1:1 (attacker : defender), with 1 μL of the resulting mixtures plated onto 1.5% LB agar pads on glass microscope slides (agar supplemented with SYTOX blue live/dead stain at 1:10^4^-fold dilution). A glass coverslip was then placed on top of each pad, and the sandwich imaged using a Nikon Ti-E inverted motorized microscope with Perfect Focus System and Plan Apo 100× Oil Ph3 DM (NA 1.4) objective. We used a Spectra X light engine (Lumencor), ET-GFP (Chroma 49002), and ET-mCherry (Chroma 49008) filter sets to excite and filter fluorescence, and a sCMOS camera pco.edge 4.2 (PCO, Germany) (pixel size 65 nm) and VisiView software (Visitron Systems, Germany) to record images. Temperature control was set at 30°c, and humidity was adjusted to 95% by an Okolab T-unit (Okolab), as in a previous publication^62^.

#### Image analysis

To analyse the dynamics of *E. coli* killing against different attackers (Tae1 and Tle1), we manually counted *E. coli* cell deaths in each movie, using the onset of SYTOX blue signal as the marker for cell death (corroborated by other toxin-specific features: sudden lysis and cell blebbing vs. Tae1; cell shrinkage followed by reinflation vs. Tle1^49^). Cell death counting was achieved by dividing each field of view into a 4×4 array of 1000 μm^2^ square regions centred on the image centroid. This left a narrow border between the array and the image edge, such that the neighbours of every E. coli cell within the array could be entirely seen. Cell deaths occurring within each movie frame were tabulated and normalised by the number of *E. coli* cells with at least one ADP1 cell contact, measured at the mid-point (frame 30, 15 minutes) of the movie. This gave a measure of cell death as a function of time, normalised to account for different levels of attacker exposure (varying naturally across experiments). Tabulated killing data were then analysed using R. We performed microscopy in full biological triplicate on at least two separate days. To minimise operator bias during cell death counting, the operator was blinded as to strain identity until data had been collected.

### Resistance costs

#### Costs in monoculture

We performed platereader growth assays in 96-well flat-bottomed microtitre plates (Corning), using a 200 μL working volume and sealing plates with a gas-permeable “Breathe-Easy” membrane (Sigma-Aldritch) to prevent well-to-well contamination. To minimise evaporation effects, replicates were arranged centrally on plates with a buffer of LB-only wells around the plate edge. Monocultures of each strain were inoculated by serially diluting overnight cultures (18 h, 37°c, 180 RPM shaking) 1:1000 (3 rounds of 10-fold dilution) into fresh LB. Plates were then transferred to a 4-bed BioTek platereader, and incubated overnight (24 h, 37°c, 500 RPM shaking), measuring OD_600_ every 20 minutes. We used 6 pseudobiological replicates: single overnight cultures inoculated from clonal cryostocks were diluted and divided to inoculate 6 separate test cultures. For reference, we included both blank (LB-only), eGFP-Anc and mCherry-Anc controls. To calculate Malthusian parameters (k_max_) for each strain, we used the equation

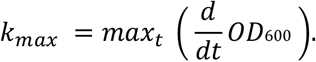

#### Costs in co-culture

Co-culture growth assays were performed in parallel to monoculture assays using flow cytometry. We mixed each eGFP-labelled overnight culture from the monoculture assays 1:1 with the mCherry-labelled ancestral strain (100 μL + 100 μL) and diluted 1:1000 into 200 μL fresh LB arranged in 96-well plates, as above (6 pseudobiological replicates). Undiluted mixes (“t_start_” sample) were held at 5°c for 30 minutes to arrest growth before being fixed (30-minute incubation in 4% paraformaldehyde solution at room temperature, ∼19 degrees) and washed (3x in 200 μL sterile-filtered dulbecco’s phosphate buffered saline, “dPBS”). Inoculated plates were then incubated alongside monoculture plates (BioTek platereader, 24 h 500 RPM shaking at 37 degrees, plates sealed with BreathEasy seals). Following incubation, we repeated our fixation procedure to prepare “t_end_” samples, which were then held at 5 degrees for 120 h prior to resuspension and flow cytometry.

Fixed t_start_ and t_end_ samples were then used to determine the initial and final ratios of eGFP-labelled mutant and mCherry-labelled ancestral strains, using a Beckman Coulter CytoFlex S flow cytometer (5 s sample mixing before measurement, 17 μL/mL sample flow rate, 5 s backwashing to clean probe after reading), exciting fluorescence with 488 nm (eGFP) and 561 nm (mCherry) lasers. Fluorescence emissions were detected at 610 nm (ECD, mCherry) and at 525 nm (FITC, eGFP), with bandpasses of 20 and 40 nm respectively. To distinguish between bacterial-sized objects and small debris, we used cell-free controls (“LB”) to determine global side- and front-scatter thresholds (SSC: 10^3^; FSC: 10^3^; Fig. S5A, B). After size thresholding, we used single-strain controls (eGFP-Anc and mCherry-Anc) to determine fluorescence thresholds to distinguish between green-and red-fluorescent bacteria (green/FITC: 10^3.05^; red/ECD: 10^2.90^; Fig. S5C, D). As additional quality control, we verified these thresholds resulted in i) <5% bacterial events neither green nor red (suggesting non-fluorescent contaminants or immature fluorophores) and ii) <5% events that were both green and red (suggesting clumping of cells; Fig. S5E, F). To calculate the competitive index of each strain, we used the equation

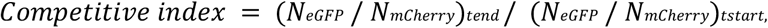

where N_eGFP_ and N_mCherry_ are respectively the numbers of green- and red-gated bacterial cells.

### Resistance from gene loss-of-function

#### Resistance assays with Keio strains

We tested single-gene knockout strains from the *E. coli* Keio collection^43^ for resistance to Tae1 amidase or Tle1 lipase toxins, using a standard CFU-based competition assay^34^. Exponential phase cultures of *E. coli* (defender) and *A. baylyi* (attacker) strains were prepared as above in full biological triplicate, washing and normalising cell densities to OD_600_ = 0.1. We prepared 1:1 mixtures of normalised cultures, plating 50 μL of each mixture onto 13mm MCE discs placed on pre-dried LB agar plates, and leaving unlidded plates in a laminar flow hood until droplets had dried (∼30 minutes). We then incubated inverted plates for 3h at 30 degrees before recovering cells, transferring each MCE disc into 1 mL sterile dPBS, and vortexing for 10 seconds. To enumerate *E. coli* and *A. baylyi* survival, we performed serial dilutions on recovered cell mixtures (7 rounds of 10-fold dilution in dPBS), plating 5 μL of each dilution in technical triplicate on pre-dried 120 mm square vented petri dishes filled with 40 mL 1.5% w/v LB agar plates, supplemented either with 100 μm/mL Streptomycin sulfate (selecting for *A. baylyi*) or 25 μm/mL Kanamycin sulfate (selecting for *E. coli)*. We incubated CFU plates at 30 °C (*A. baylyi* selection) or 37 °C (*E. coli* selection) overnight (∼16 h) until individual colonies became visible. We counted the number of CFUs at the highest dilution at which at least 5 CFUs were visible, and then multiplied this number by the dilution factor. To account for plate-to-plate variation in CFU count from a given recovered cell suspension, we took the mean count over 3 technical replicate platings for each competition replicate.

### Data Processing and Statistics

Data were analysed and visualized using Rstudio (Version 2024.12.0+467), running R 4.3.2 (2023-10-31; “eyeholes”), and using the packages readxl, ggplot2, cowplot, dplyrk, tidyr and the multcomp statistics library. Colorimetric data were processed using Matlab R2024b (V.24.2.0.2833386). Timelapse microscopy data were analysed using FIJI (ImageJ2 V.2.14.0/1.54f) open source image processing software, using the Bio-Formats Java library. Cytometry data was collected and gated using CytoFLEX CytExpert software before being exported and processed in R using the flowCore package. Statistical tests were performed with using linear modelling with ANOVA (via R’s inbuilt lm method) to compare *E. coli* mutants and ancestors, using biological replicate as a fixed factor, and using Dunnett’s test for multiple comparisons (multcomp methods: glht, mcp).

## Acknowledgements

We are grateful to Claudia Igler, Andrew Preston, Taoran Fu, Norman van Rhijn and Gavin Thomas for helpful comments during manuscript drafting, and to Rok Krašovec for use of Keio collection strains. We would like to thank Matthew Thomas and Rosanna Wright for training and technical assistance with flow cytometry. This work was supported by the National Center of Competence in Research (“AntiResist”), funded by the Swiss National Science Foundation (51NF40_180541) to MB and ATA. RT is funded by a University of Manchester Dame Katheleen Ollerenshaw Fellowship. WPJS is funded by a Wellcome Trust Sir Henry Wellcome PostDoctoral Fellowship (222795/Z/21/Z) with additional funding from a Wellcome Trust University of Manchester Translational Partnership Award (222061/Z/20/Z).

## Author Contributions

Conceptualization, Investigation, Funding acquisition: WPJS, MB and MAB. WPJS and ATA WPJS and ATA performed experiments. Methodology and data analysis by WPJS, ATA and RT. WPJS and RT wrote software to analyse the data. WPJS drafted the manuscript; WPJS, ATA, RT, MB and MAB reviewed and edited the manuscript.

## References

1. Hibbing, M. E., Fuqua, C., Parsek, M. R. & Peterson, S. B. Bacterial competition: surviving and thriving in the microbial jungle. Nat Rev Microbiol 8, 15–25 (2010).

2. Granato, E. T., Meiller-Legrand, T. A. & Foster, K. R. The Evolution and Ecology of Bacterial Warfare. Current Biology 29, R521–R537 (2019).

3. Peterson, S. B., Bertolli, S. K. & Mougous, J. D. The Central Role of Interbacterial Antagonism in Bacterial Life. Current Biology 30, R1203–R1214 (2020).

4. Stubbusch, A. K. M. et al. Antagonism as a foraging strategy in microbial communities. Science 388, 1214–1217 (2025).

5. Ross, B. D. et al. Human gut bacteria contain acquired interbacterial defence systems. Nature 575, 224–228 (2019).

6. Haas, A. L. et al. Iron bioavailability regulates Pseudomonas aeruginosa interspecies interactions through type VI secretion expression. Cell Rep 42, 112270 (2023).

7. Speare, L. et al. Bacterial symbionts use a type VI secretion system to eliminate competitors in their natural host. Proceedings of the National Academy of Sciences 115, E8528–E8537 (2018).

8. Sana, T. G. et al. Salmonella Typhimurium utilizes a T6SS-mediated antibacterial weapon to establish in the host gut. Proceedings of the National Academy of Sciences 113, E5044– E5051 (2016).

9. Sana, T. G., Lugo, K. A. & Monack, D. M. T6SS: The bacterial ‘fight club’ in the host gut. PLOS Pathogens 13, e1006325 (2017).

10. Toska, J., Ho, B. T. & Mekalanos, J. J. Exopolysaccharide protects Vibrio cholerae from exogenous attacks by the type 6 secretion system. Proceedings of the National Academy of Sciences 115, 7997–8002 (2018).

11. Hersch, S. J. et al. Envelope stress responses defend against type six secretion system attacks independently of immunity proteins. Nat Microbiol 5, 706–714 (2020).

12. Hersch, S. J., Manera, K. & Dong, T. G. Defending against the Type Six Secretion System: beyond Immunity Genes. Cell Reports 33, 108259 (2020).

13. Robitaille, S., Trus, E. & Ross, B. D. Bacterial Defense against the Type VI Secretion System. Trends Microbiol 29, 187–190 (2021).

14. Granato, E. T., Smith, W. P. J. & Foster, K. R. Collective protection against the type VI secretion system in bacteria. The ISME Journal 17, 1052–1062 (2023).

15. Smith, W. P. J., Wucher, B. R., Nadell, C. D. & Foster, K. R. Bacterial defences: mechanisms, evolution and antimicrobial resistance. Nat Rev Microbiol 21, 519–534 (2023).

16. Trotta, K. L. et al. Lipopolysaccharide transport regulates bacterial sensitivity to a cell wall-degrading intermicrobial toxin. PLOS Pathogens 19, e1011454 (2023).

17. MacGillivray, K. A. et al. Trade-offs constrain adaptive pathways to type VI secretion system survival. iScience 26, 108332 (2023).

18. Kennedy, N. W. & Comstock, L. E. Mechanisms of bacterial immunity, protection, and survival during interbacterial warfare. Cell Host & Microbe 32, 794–803 (2024).

19. Dessartine, M. M., Kosta, A., Doan, T., Cascales, É. & Côté, J.-P. Type 1 fimbriae-mediated collective protection against type 6 secretion system attacks. mBio 15, e02553–23 (2024).

20. Flaugnatti, N., Bader, L., Croisier-Coeytaux, M. & Blokesch, M. Capsular polysaccharide restrains type VI secretion in Acinetobacter baumannii. Elife 14, e101032 (2025).

21. Tejada-Arranz, A. et al. Mechanisms of Pseudomonas aeruginosa resistance to type VI secretion system attacks. Nat Commun 16, 10744 (2025).

22. Boyer, F., Fichant, G., Berthod, J., Vandenbrouck, Y. & Attree, I. Dissecting the bacterial type VI secretion system by a genome wide in silico analysis: what can be learned from available microbial genomic resources? BMC Genomics 10, 104 (2009).

23. Wexler, A. G. et al. Human symbionts inject and neutralize antibacterial toxins to persist in the gut. Proceedings of the National Academy of Sciences 113, 3639–3644 (2016).

24. Steele, M. I. & Moran, N. A. Evolution of Interbacterial Antagonism in Bee Gut Microbiota Reflects Host and Symbiont Diversification. mSystems 6, 10.1128/msystems.00063-21 (2021).

25. Williams, D. J. et al. Competitive behaviors in Serratia marcescens are coordinately regulated by a lifestyle switch frequently inactivated in the clinical environment. Cell Host & Microbe 33, 252-266.e5 (2025).

26. Bernal, P., Allsopp, L. P., Filloux, A. & Llamas, M. A. The Pseudomonas putida T6SS is a plant warden against phytopathogens. ISME J 11, 972–987 (2017).

27. Bernal, P., Llamas, M. A. & Filloux, A. Type VI secretion systems in plant-associated bacteria. Environmental Microbiology 20, 1–15 (2018).

28. Coulthurst, S. The Type VI secretion system: a versatile bacterial weapon. Microbiology 165, 503–515 (2019).

29. Russell, A. B., Peterson, S. B. & Mougous, J. D. Type VI secretion effectors: poisons with a purpose. Nat Rev Microbiol 12, 137–148 (2014).

30. LaCourse, K. D. et al. Conditional toxicity and synergy drive diversity among antibacterial effectors. Nat Microbiol 3, 440–446 (2018).

31. Zhang, J. et al. SecReT6 update: a comprehensive resource of bacterial Type VI Secretion Systems. SCLS 66, 626–634 (2022).

32. Smith, W. P. J. et al. The evolution of the type VI secretion system as a disintegration weapon. PLOS Biology 18, e3000720 (2020).

33. Booth, S. C., Smith, W. P. J. & Foster, K. R. The evolution of short- and long-range weapons for bacterial competition. Nat Ecol Evol 7, 2080–2091 (2023).

34. Smith, W. P. J. et al. Multiplicity of type 6 secretion system toxins limits the evolution of resistance. Proceedings of the National Academy of Sciences 122, e2416700122 (2025).

35. Ting, S.-Y. et al. Discovery of coordinately regulated pathways that provide innate protection against interbacterial antagonism. eLife 11, e74658.

36. Kirchberger, P. C., Unterweger, D., Provenzano, D., Pukatzki, S. & Boucher, Y. Sequential displacement of Type VI Secretion System effector genes leads to evolution of diverse immunity gene arrays in Vibrio cholerae. Sci Rep 7, 45133 (2017).

37. Azhieh, A. et al. Rapidly evolving orphan immunity genes protect human gut bacteria from intoxication by the type VI secretion system. 2025.05.03.651265 Preprint at 10.1101/2025.05.03.651265 (2025).

38. Lin, H.-H. et al. A High-Throughput Interbacterial Competition Screen Identifies ClpAP in Enhancing Recipient Susceptibility to Type VI Secretion System-Mediated Attack by Agrobacterium tumefaciens. Front Microbiol 10, 3077 (2019).

39. Le, N.-H. et al. Peptidoglycan editing provides immunity to Acinetobacter baumannii during bacterial warfare. Sci Adv 6, eabb5614 (2020).

40. Lories, B. et al. Biofilm Bacteria Use Stress Responses to Detect and Respond to Competitors. Curr Biol 30, 1231-1244.e4 (2020).

41. Borenstein, D. B., Ringel, P., Basler, M. & Wingreen, N. S. Established Microbial Colonies Can Survive Type VI Secretion Assault. PLOS Computational Biology 11, e1004520 (2015).

42. Custodio, R. et al. Type VI secretion system killing by commensal Neisseria is influenced by expression of type four pili. eLife 10, e63755 (2021).

43. Baba, T. et al. Construction of Escherichia coli K-12 in-frame, single-gene knockout mutants: the Keio collection. Mol Syst Biol 2, 2006.0008 (2006).

44. Bottery, M. J., Wood, A. J. & Brockhurst, M. A. Adaptive modulation of antibiotic resistance through intragenomic coevolution. Nat Ecol Evol 1, 1364–1369 (2017).

45. Cooper, T. F., Rozen, D. E. & Lenski, R. E. Parallel changes in gene expression after 20,000 generations of evolution in Escherichia coli. Proceedings of the National Academy of Sciences 100, 1072–1077 (2003).

46. Karp, P. D. et al. The EcoCyc Database (2023). EcoSal Plus 11, eesp-0002-2023 (2023).

47. The UniProt Consortium. UniProt: the Universal Protein Knowledgebase in 2025. Nucleic Acids Res 53, D609–D617 (2025).

48. Hachani, A., Lossi, N. S. & Filloux, A. A visual assay to monitor T6SS-mediated bacterial competition. J Vis Exp e50103 (2013) doi:10.3791/50103.

49. Ringel, P. D., Hu, D. & Basler, M. The Role of Type VI Secretion System Effectors in Target Cell Lysis and Subsequent Horizontal Gene Transfer. Cell Rep 21, 3927–3940 (2017).

50. Meier-Dieter, U., Barr, K., Starman, R., Hatch, L. & Rick, P. D. Nucleotide sequence of the Escherichia coli rfe gene involved in the synthesis of enterobacterial common antigen. Molecular cloning of the rfe-rff gene cluster. Journal of Biological Chemistry 267, 746–753 (1992).

51. Yanni, D. et al. Life in the coffee-ring: how evaporation-driven density gradients dictate the outcome of inter-bacterial competition. Preprint at 10.48550/arXiv.1707.03472 (2017).

52. Choi, U., Park, S. H., Lee, H. B., Son, J. E. & Lee, C.-R. Coordinated and Distinct Roles of Peptidoglycan Carboxypeptidases DacC and DacA in Cell Growth and Shape Maintenance under Stress Conditions. Microbiology Spectrum 11, e00014–23 (2023).

53. Hanga, K.-S. N., Brockhurst, M. A. & Bottery, M. J. Chromosomal resistance mutations facilitate acquisition of multidrug-resistant plasmids in Escherichia coli. Microbiology 171, 001599 (2025).

54. Zhu, K., Zhang, Y.-M. & Rock, C. O. Transcriptional regulation of membrane lipid homeostasis in Escherichia coli. J Biol Chem 284, 34880–34888 (2009).

55. Hersch, S. J., Sejuty, R. T., Manera, K. & Dong, T. G. High throughput identification of genes conferring resistance or sensitivity to toxic effectors delivered by the type VI secretion system. 2021.10.06.463450 Preprint at 10.1101/2021.10.06.463450 (2021).

56. Crisan, C. V. et al. Glucose confers protection to Escherichia coli against contact killing by Vibrio cholerae. Sci Rep 11, 2935 (2021).

57. Tokishita, S., Kojima, A. & Mizuno, T. Transmembrane signal transduction and osmoregulation in Escherichia coli: functional importance of the transmembrane regions of membrane-located protein kinase, EnvZ. J Biochem 111, 707–713 (1992).

58. Ferrières, Lionel., Aslam, S. N., Cooper, R. M. & Clarke, D. J. The yjbEFGH locus in Escherichia coli K-12 is an operon encoding proteins involved in exopolysaccharide production. Microbiology 153, 1070–1080 (2007).

59. Kitano, H. Towards a theory of biological robustness. Mol Syst Biol 3, 137 (2007).

60. Whitacre, J. M. Biological Robustness: Paradigms, Mechanisms, and Systems Principles. Front Genet 3, 67 (2012).

61. Deatherage, D. E. & Barrick, J. E. Identification of mutations in laboratory-evolved microbes from next-generation sequencing data using breseq. Methods Mol Biol 1151, 165–188 (2014).

62. George, M., Narayanan, S., Tejada-Arranz, A., Plack, A. & Basler, M. Initiation of H1-T6SS dueling between Pseudomonas aeruginosa. mBio 15, e00355–24 (2024).

