## Supplementary Information for "Evolved resistance against the type 6 secretion system is toxin specific"

### Author information

William Smith[1] * †(ORCID),
Alejandro Tejada Arranz[2] *(ORCID),
Raveen Tank[1](ORCID),
Marek Basler[2] (ORCID),
Michael Brockhurst[1] †(ORCID).

1. University of Manchester, Manchester, United Kingdom.
2. University of Basel, Basel, Switzerland

†: corresponding author:


*=equal contribution

### Supplementary Tables

**Table S1: Bacterial strains used in this study**

| Name | Species / strain | Genotype | Comments | Origin |
|---|---|---|---|---|
| *Ab* WT | *A. baylyi* ADP1 | *rpsL-K88R* | Parental ADP1 T6SS attacker strain; streptomycin resistance marker | Ringel *et al.* [1] |
| *Ab* ΔE | *A. baylyi* ADP1 | *rpsL-K88R clpV-mCherry2 Δaciad0053 Δaciad0168 Δaciad1790 Δaciad3114 Δaciad3425* | Attacker strain lacking all known T6SS effectors | Ringel *et al.* [1] |
| *Ab* Tle1 | *A. baylyi* ADP1 | *rpsL-K88R clpV-mCherry2 Δaciad0053 Δaciad0168 Δaciad1790 Δaciad3114* | Attacker strain carrying only effector ACIAD3425 (Tle1) | Ringel *et al.* [1] |
| *Ab* Tae1 | *A. baylyi* ADP1 | *rpsL-K88R clpV-mCherry2 Δaciad0053 Δaciad1790 Δaciad3114 Δaciad3425* | Strain carrying only effector ACIAD0168 (Tae1) | Ringel *et al.* [1] |
| *Ab* Tae1+Tle1 | *A. baylyi* ADP1 | *rpsL-K88R clpV-mCherry2 Δaciad0053 Δaciad1790 Δaciad3114* | Strain carrying only effectors Tae1 and Tle1 | Ringel *et al.* [1] |
| *Ab Δhcp* | *A. baylyi* ADP1 | *rpsL-K88R, Δhcp vipA-sfGFP clpVmCherry2 ADP1 with no functional T6SS* | T6SS negative control to test *Ec* adaptation to co-culture | Ringel *et al.* [1] |
| *Vc* WT | *V. cholerae* 2740-80 | Sm$^R$, LacZ- | Alternative T6SS attacker strain; streptomycin resistance marker | Basler et al. [2] |
| *Vc* µ*vasX* | *V. cholerae* 2740-80 | Sm$^R$, LacZ- VasX-ΔA852-F867 | *Vc* with pore-forming toxin VasX inactivated via point mutation | This study |
| *Vc* µE | *V. cholerae* 2740-80 | Sm$^R$, LacZ- VgrG3-D842A VasX-ΔA852-F867 TseH-H64A TseL-D425A VipA-mCh2 | *Vc* with all known toxins inactivated via point mutations | Tejada-Arranz et al.[3] |

| | | | | |
|---|---|---|---|---|
| *Ec* Anc | *E. coli* MG1665 | *attB::eGFP kan*$^R$ | Ancestral *E. coli* strain carrying eGFP tag and kanamycin resistance marker | Smith *et al.* [4] |
| *Ec* A1 | *E. coli* MG1665 | *attB::eGFP kan*$^R$+ *mutations* | Tae1-evolved | Smith *et al.* [4] |
| *Ec* A2 | *E. coli* MG1665 | *attB::eGFP kan*$^R$+ *mutations* | Tae1-evolved | Smith *et al.* [4] |
| *Ec* A3 | *E. coli* MG1665 | *attB::eGFP kan*$^R$+ *mutations* | Tae1-evolved | Smith *et al.* [4] |
| *Ec* A4 | *E. coli* MG1665 | *attB::eGFP kan*$^R$+ *mutations* | Tae1-evolved | Smith *et al.* [4] |
| *Ec* A5 | *E. coli* MG1665 | *attB::eGFP kan*$^R$+ *mutations* | Tae1-evolved | Smith *et al.* [4] |
| *Ec* A6 | *E. coli* MG1665 | *attB::eGFP kan*$^R$+ *mutations* | Tae1-evolved | Smith *et al.* [4] |
| *Ec* A7 | *E. coli* MG1665 | *attB::eGFP kan*$^R$+ *mutations* | Tae1-evolved | Smith *et al.* [4] |
| *Ec* A8 | *E. coli* MG1665 | *attB::eGFP kan*$^R$+ *mutations* | Tae1-evolved | Smith *et al.* [4] |
| *Ec* B2 | *E. coli* MG1665 | *attB::eGFP kan*$^R$+ *mutations* | Tle1-evolved | Smith *et al.* [4] |
| *Ec* B3 | *E. coli* MG1665 | *attB::eGFP kan*$^R$+ *mutations* | Tle1-evolved | Smith *et al.* [4] |
| *Ec* B4 | *E. coli* MG1665 | *attB::eGFP kan*$^R$+ *mutations* | Tle1-evolved | Smith *et al.* [4] |
| *Ec* B6 | *E. coli* MG1665 | *attB::eGFP kan*$^R$+ *mutations* | Tle1-evolved | Smith *et al.* [4] |
| *Ec* B8 | *E. coli* MG1665 | *attB::eGFP kan*$^R$+ *mutations* | Tle1-evolved | Smith *et al.* [4] |
| *Ec* C6 | *E. coli* MG1665 | *attB::eGFP kan*$^R$+ *mutations* | Tae1+Tle1-evolved | Smith *et al.* [4] |
| *Ec ΔlacY* | *E. coli* BW25113 | *ΔlacY::kan*$^R$ | LacY knockout, control for Keio competitions | Baba *et al.* [5] |
| *Ec ΔwecA* | *E. coli* BW25113 | *ΔwecA::kan*$^R$ | WecA (rfe) knockout | Baba *et al.* [5] |
| *Ec ΔdctA* | *E. coli* BW25113 | *ΔdctA::kan*$^R$ | DctA knockout | Baba *et al.* [5] |
| *Ec ΔenvZ* | *E. coli* BW25113 | *ΔenvZ::kan*$^R$ | EnvZ knockout | Baba *et al.* [5] |
| *Ec ΔnlpI* | *E. coli* BW25113 | *ΔnlpI::kan*$^R$ | NlpI knockout | Baba *et al.* [5] |
| *Ec ΔsstT* | *E. coli* BW25113 | *ΔsstT::kan*$^R$ | SstT knockout | Baba *et al.* [5] |
| *Ec ΔplsY* | *E. coli* BW25113 | *ΔplsY::kan*$^R$ | PlsY knockout | Baba *et al.* [5] |
| *Ec ΔdacA* | *E. coli* BW25113 | *ΔdacA::kan*$^R$ | DacA knockout | Baba *et al.* [5] |
| *Ec ΔdpaA* | *E. coli* BW25113 | *ΔdpaA::kan*$^R$ | DpaA knockout | Baba *et al.* [5] |
| *Ec ΔyobF* | *E. coli* BW25113 | *ΔyobF::kan*$^R$ | YobF knockout | Baba *et al.* [5] |

### Supplementary Figures

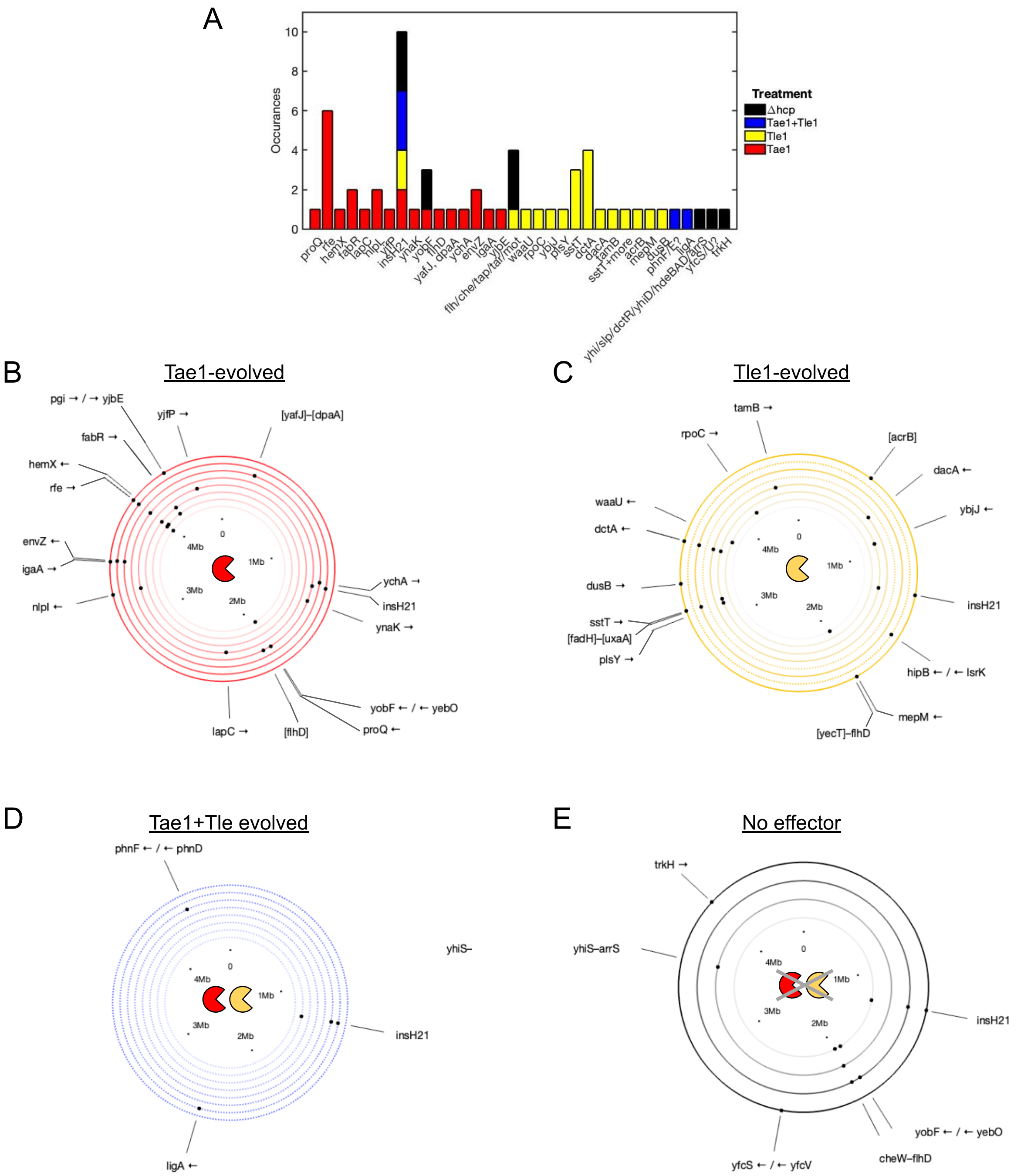


**Fig. S1: T6SS-evolved *E. coli* show extensive parallel evolution within effector treatments. (A)** Summary of mutation instances in end-point *E. coli* isolates grouped by the T6SS attacker treatment group in which they occurred (legend). **(B)** Ring diagram summarising mutations arising in end-point clonal isolates evolved against amidase effector Tae1. Rings represent independent lineages A1 (innermost) to A8 (outermost), with dots representing mutations at specified locations on the chromosome (annotated according to gene(s) affected). **(C)** as (B) but for lipase Tle1-evolved lineages. Dashed rings correspond to populations that went extinct before the final (30$^{th}$) transfer. **(D)** As (C) but for lineages evolved against both Tae1 and Tle1. **(E)** as (C) but for four lineages evolved against a T6SS− (*Δhcp*) control strain.

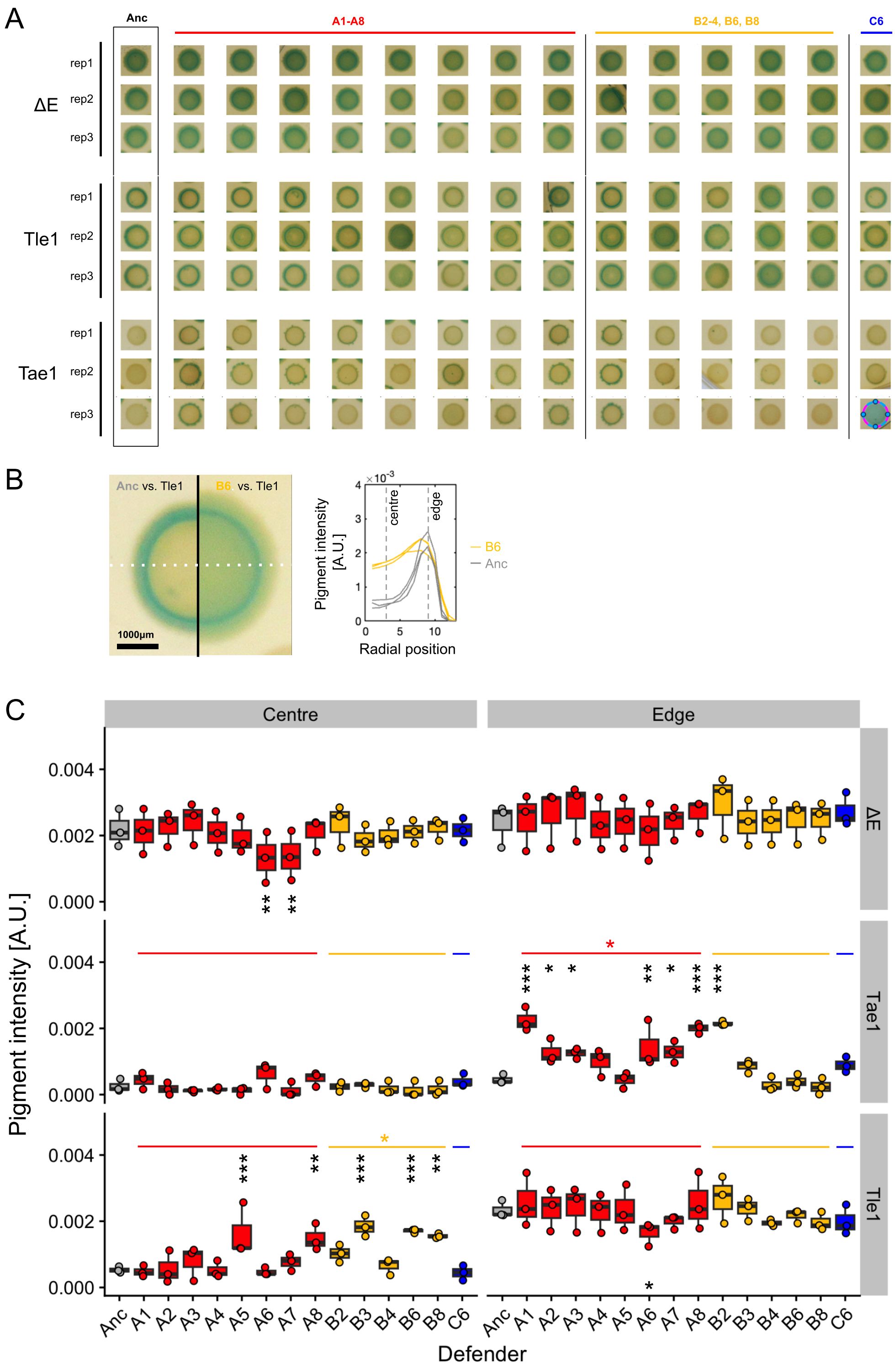


**Fig. S2: Colorimetric competition assays reveal toxin-specific resistance at colony edge or centre, but rule out aggregation or edge escape as resistance mechanisms. (A)** Complete dataset of XGal-agar colony images taken after 16h incubation. Each column corresponds to a different *E. coli* isolate (A1-8: Tae1-evolved; B2-4/6/8: Tle1-evolved; C6: Tae1+Tle1-evolved); rows show biological (rep)licate cocultures with ΔE (effectorless), Tle1- or Tae1-armed *A. baylyi*. For each treatment, replicate 1/3 is shown in Fig. 2. (**B)** Zoomed images of Anc(estral) and B6 *E. coli* in co-culture with Tle1-armed *A. baylyi* (left) shown alongside radial XGal pigmentation traces (right), highlighting increased pigmentation in colony centre (evaluated at 20-30% colony radius) but not at colony edge (90-100% colony radius). **(C)** Pigmentation intensities at colony centres vs. edges for *E. coli* defenders (horizontal axes) confronted with different *A. baylyi* attackers (rows). Black asterisks mark individual isolates with significantly altered pigmentation at colony centre or edges compared with the ancestral strain (significance codes: p<0.1, * p < 0.05, ** p<0.01, *** p<0.001). Tests performed using linear regression (R: “lm” method using Dunnett’s contrast for multiple comparisons). Coloured asterisks mark evolutionary treatment groups with significantly altered pigmentation cf. the ancestral strain in each location. N=3 biological replicates per Defender / Attacker combination.

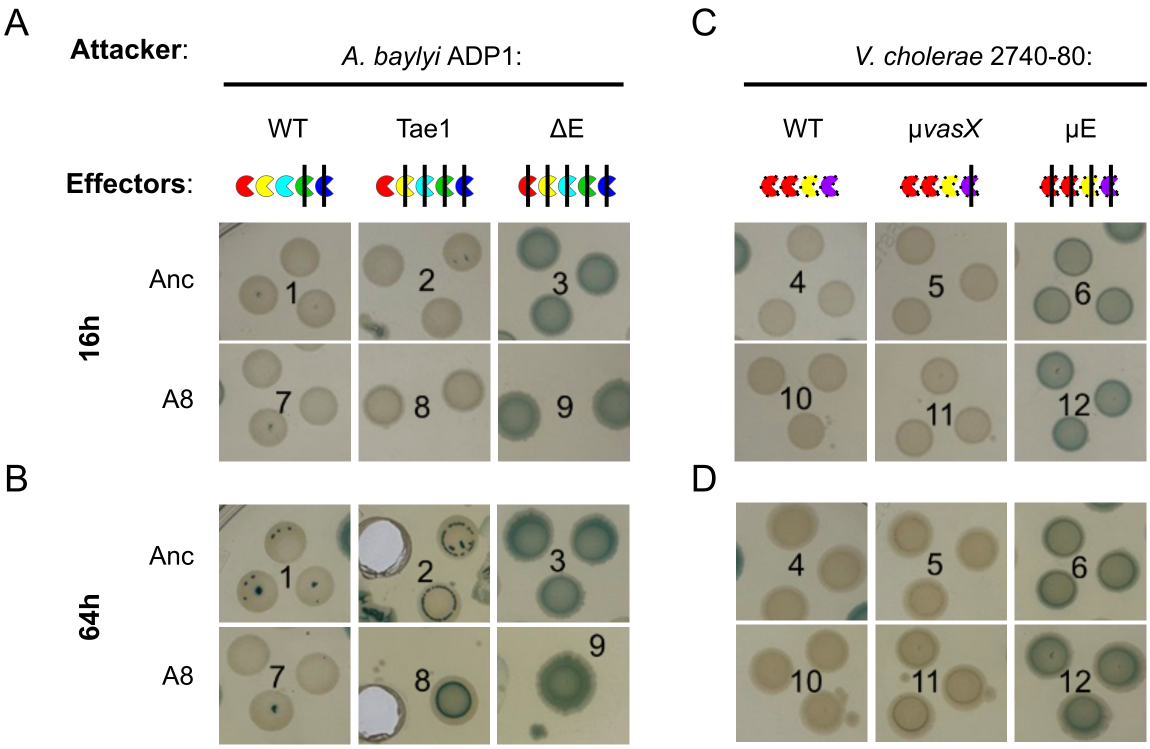


**Fig. S3: Further colorimetric tests of cross-resistance to alterate T6SS effectors. (A)** XGal competition assay comparing survival Anc(estral) and A8 (Tae1-evolved) *E. coli* defenders in 16h co-culture with T6SS+ ADP1 attackers armed with 3 (WT: Tae1+Tle1+Tse2), 1 (Tae1) or 0 (ΔE) effectors. **(B)** Same colonies as in (A) but after 64h incubation. **(C)** as (A) but against *V. cholerae* T6SS+ attackers armed with four (WT = VgrG3+TseH+TseL+VasX), three (μ*vasX* = VgrG3+TseH+TseL), or zero (μE) active effectors. **(D)** as (C) but imaged after 64h. All competitions carried out in full biological triplicate; each panel shows 3 independent co-cultures with the exception of "8" (A8 vs. Tae1). Note that some colonies were excised for further analysis after 16h, and so are not visible in (B, D).

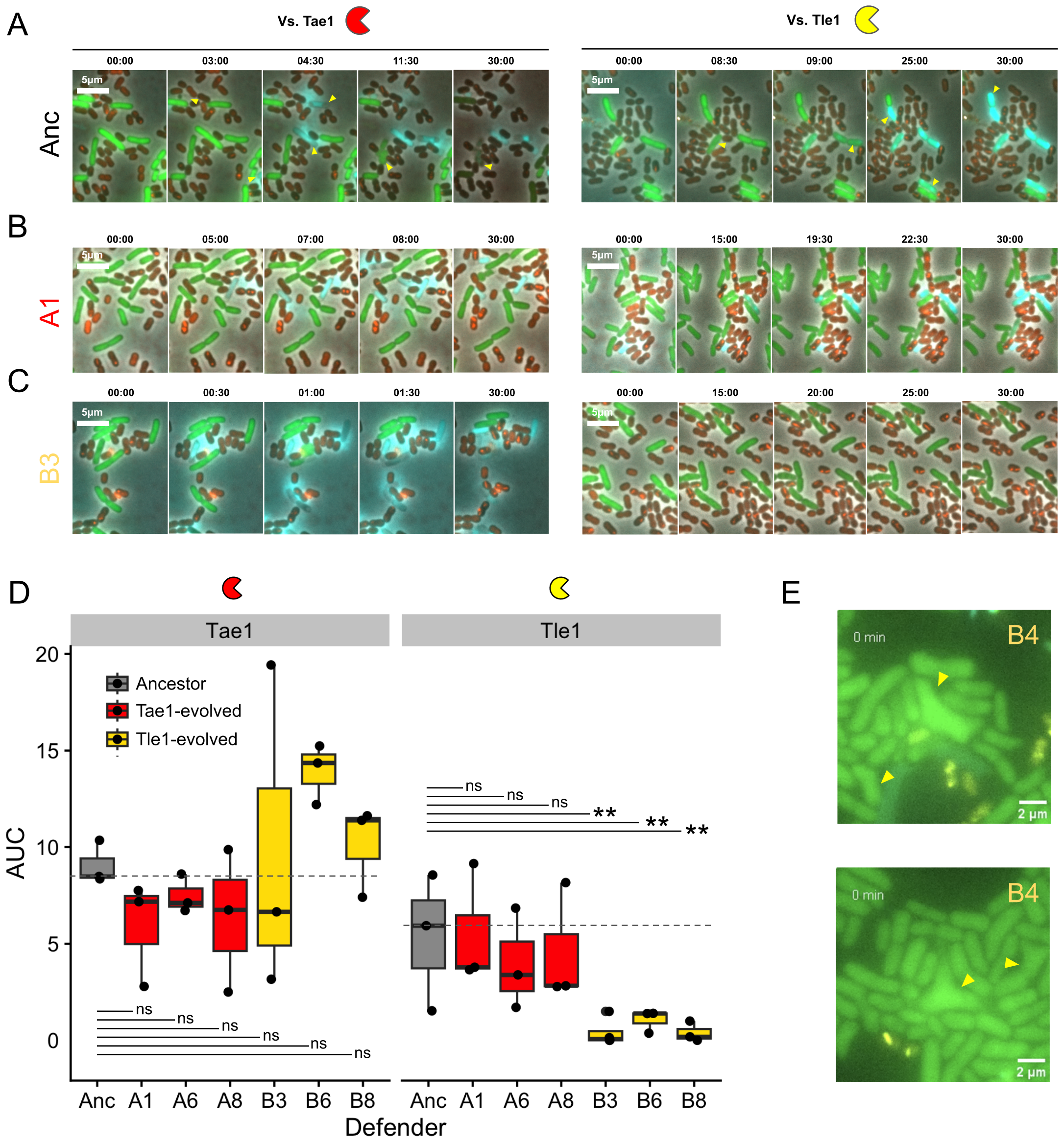


**Fig. S4: Comparisons of single-cell phenotypes and killing dynamics. (A):** Representative snapshots from 30m timelapse microscopy, showing the Anc(estral) *E. coli* strain's responses to ADP1 attackers armed with Tae1 amidase (left column; fast lysis phenotype) or Tle1 lipase (right column, slow permeabilization phenotype). Image timestamps and scale bars as shown; eGFP+ *E. coli* cells, T6SS foci (vipA::mCherry fusion) and SYTOX permeability stain are respectively coloured green, red and cyan. Yellow arrows highlight individual cell death death events. **(B):** as A, but with Tae1-evolved *E. coli* isolate A1. **(C):** as A, but with Tae1-evolved *E. coli* isolate B3. **(D)** Areas under normalised *E. coli* killing curves shown in Fig. 3 (AUC), plotted for cocultures with amidase- (Tae1, left) and lipase-armed *A. baylyi* bacteria. Boxplots are coloured according to the treatment used to evolve each Defender strain (Red: Tae1; Yellow, Tle1). Statistical tests performed by comparing the ancestral killing curve (Grey) with each strain using a linear model (R: lm(AUC ~ Defender + Replicate), incorporating Dunnett's test for multiple comparisons. N=3 biological replicates conducted on different days per Defender / Attacker combination. Tests of significance: ns = non-significant ($p > 0.05$); ** = $p < 0.01$). **(E)** examples of *E. coli* isolate B4 cells showing abnormal and branched morphologies (yellow arrows).

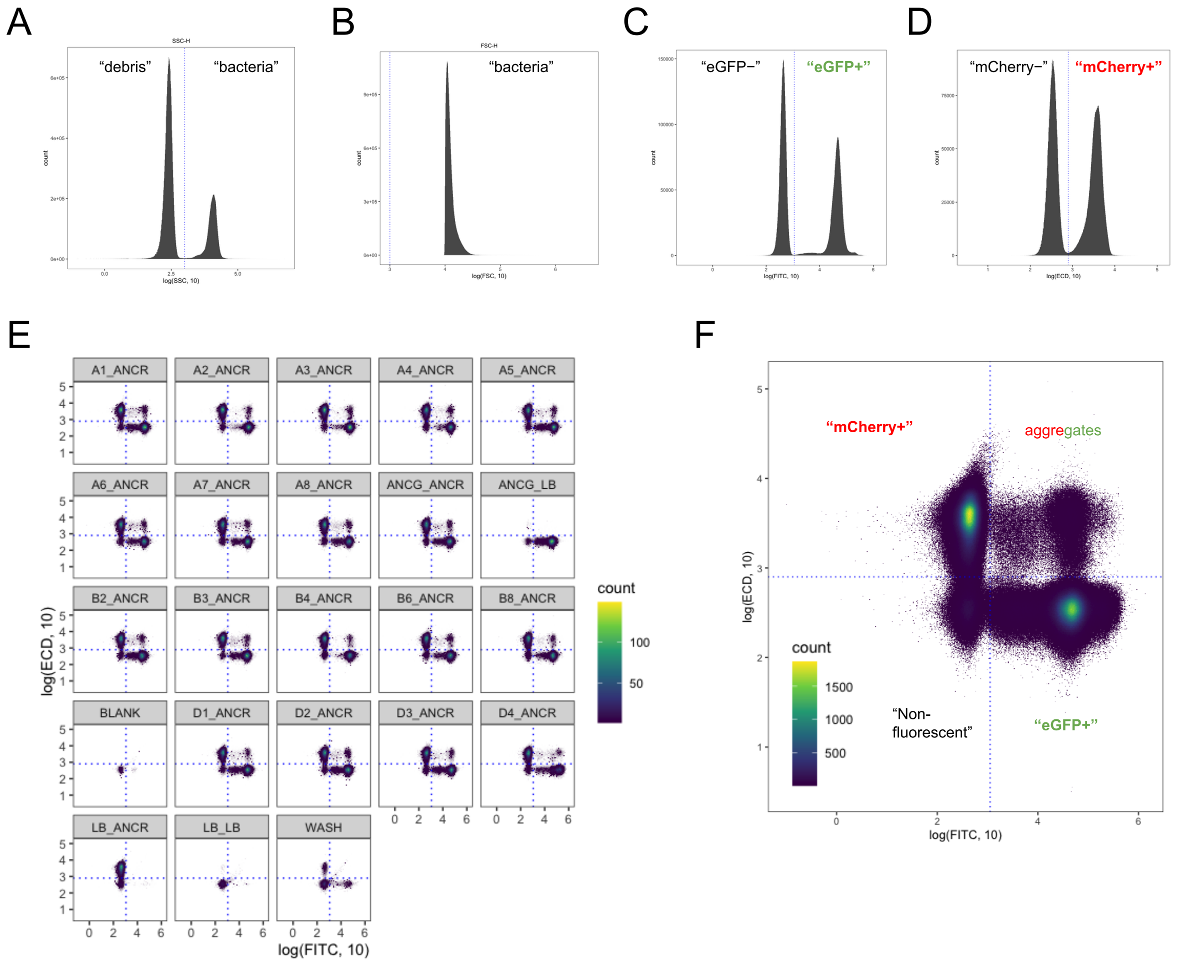


**Fig. S5: particle scatter and fluorescence gating strategy for cytometric fitness assay. (A)** Distribution of size scatter signal (SSC) aggregated across all samples and timepoints, showing SSC threshold used to exclude non-bacterial debris (“bacteria” defined as SSC>$10^{3}$). **(B)** Front scatter (FSC) distribution, not used for gating. **(C)** Distribution of green fluorescence intensity (FITC) for bacterial-sized objects gated via A, showing FITC threshold (“eGFP+” defined as FITC>$10^{3.05}$). **(D)** As C) but for red fluorescence (ECD), showing threshold (“mCherry+” defined as ECD>$10^{2.90}$). **(E)** Quadrant plots showing event density against FITC and ECD signal for all samples in competition assay, validating gating strategy (note absence of red fluorescence in green-only ANCG_LB sample and vice versa LB_ANCR. **(F)** Aggregate quadrant plot for all samples in (E).

### Supplementary References


1. Ringel, P. D., Hu, D. & Basler, M. The Role of Type VI Secretion System Effectors in Target Cell Lysis and Subsequent Horizontal Gene Transfer. *Cell Rep* **21**, 3927–3940 (2017).
2. Basler, M., Pilhofer, M., Henderson, G. P., Jensen, G. J. & Mekalanos, J. J. Type VI secretion requires a dynamic contractile phage tail-like structure. *Nature* **483**, 182–186 (2012).
3. Tejada-Arranz, A. *et al.* Mechanisms of Pseudomonas aeruginosa resistance to Type VI Secretion System attacks. 2024.10.26.620397 Preprint at https://doi.org/10.1101/2024.10.26.620397 (2025).
4. Smith, W. P. J. *et al.* Multiplicity of type 6 secretion system toxins limits the evolution of resistance. *Proceedings of the National Academy of Sciences* **122**, e2416700122 (2025).
5. Baba, T. *et al.* Construction of Escherichia coli K-12 in-frame, single-gene knockout mutants: the Keio collection. *Mol Syst Biol* **2**, 2006.0008 (2006).
